# Electrochemical Lateral Flow Assay with Linked Analytics for the Surveillance of Cassava Brown Streak Disease in East Africa

**DOI:** 10.1101/2025.10.13.682040

**Authors:** José M. R. Flauzino, Abdulkadir Sanli, Rudolph R. Shirima, Yuanjun Cai, Tinghao Hu, Leyang Li, Taliya Weinstein, Selin Olenik, Abdülkadir Gümüşçü, Laura Gonzalez-Macia, George Mahuku, James Legg, Anthony E.G. Cass, Firat Güder

## Abstract

Cassava brown streak disease (CBSD) severely threatens food security in East Africa and the livelihoods of hundreds of millions globally. Effective control requires large-scale surveillance in resource-limited settings, which is currently lacking. We present ELLA (<u>E</u>lectrochemical <u>L</u>ateral Flow Assay with <u>L</u>inked <u>A</u>nalytics), a portable, battery-free, low-cost digital diagnostic platform integrating lateral flow assays with near-field communication and electrochemical readouts for cloud-based data storage and analytics. Validated through extensive field trials in East Africa and smaller studies in Brazil, ELLA achieved 89% agreement with RT-qPCR and 95% with ELISA, often surpassing ELISA sensitivity at a material cost below US$1 per test. Leveraging ELLA’s molecular results, we trained a deep-learning model (DeepELLA) for rapid, image-based diagnosis of CBSD, enabling scalable surveillance of this and potentially emerging plant pathogens. By combining electrochemical sensing, digital connectivity, and AI-driven analytics, ELLA offers a powerful tool to strengthen plant disease monitoring and food security. Its modular design also allows adaptation to other chemical and biological targets, creating opportunities for novel datasets and new insights into plant and environmental health.

## Introduction

Cassava brown streak disease (CBSD) is a devastating viral infection caused by the Cassava brown streak virus (CBSV) and Ugandan cassava brown streak virus (UCBSV) which are positive-sense single-stranded RNA viruses belonging to the genus *Ipomovirus*.^1^ CBSD poses a major threat to food security in East Africa, where it is rapidly spreading westward. This rapid spread risks the food security of more than 800 million people worldwide who depend on cassava as a staple crop.^2^ The viruses spreads primarily via whiteflies and infected planting materials (cuttings) leading to substantial losses, including complete crop failures, in the production of cassava (*Manihot esculenta*).^3^ At later stages, the CBSD manifests itself as brown lesions on stems, yellow or necrotic vein banding on mature leaves, and dark necrotic areas within tuberous roots, degrading root quality.^4^ The observable visual symptoms, however, are typically not present in the earlier stages of infection. Some plants may also remain completely free of visual symptoms in the stems and leaves but produce lower or no yield at harvest.^5^ Furthermore, recognizing the observable visual symptoms of CBSD can be difficult even for the highly trained field scientists, hence the exact diagnosis of CBSD is usually confirmed by molecular methods.^6^

Plant pathogens often spread rapidly and exponentially^7^ which presents a series of unique challenges for molecular diagnostic testing of plant pathogens for guiding local, regional and global interventions for at least three reasons: i) Testing must be performed with technologies that produce results rapidly in the field to guide immediate local interventions. As most agricultural land in the world is also distant from urban centers, shipping samples to centralized facilities can be costly and slow delaying any immediate action; ii) Testing must be performed at scale for geographical mapping which requires low-cost technologies with geolocation capabilities. iii) For guiding regional and global interventions effectively, results must be available to national and international agencies in real-time, in a digital format to develop a data-driven action plan to address outbreaks. The urgent need for real-time, low-cost digital diagnostic tools extends beyond CBSD to a range of devastating plant diseases, including Banana bunchy top virus, Tomato yellow leaf curl virus, Maize lethal necrosis, Citrus greening, Wheat rust, and Potato late blight.^8^

The current molecular techniques for the detection of CBSD include enzyme-linked immunosorbent assay (ELISA), reverse transcription recombinase polymerase amplification (RT-RPA)^9^ and the gold standard, reverse transcription quantitative polymerase chain reaction (RT-qPCR)^10^. While these methods are sensitive and reliable, they require specialized centralized laboratory facilities (and advanced equipment). Most importantly, ELISA and RT-qPCR require several tedious sample-handling steps, which are typically performed manually by highly trained laboratory personnel.^11^ For CBSD, samples are often collected and shipped from rural regions to a small number of specialized laboratories in major cities with a turnaround time for results of several days to weeks.^12^

Lateral flow assays (LFAs) are a simple and low-cost solution to rapid testing at the point-of-use. This assay format traditionally only provides results that are qualitative – a simple yes or no with the appearance of a line in the test.^13^ Their evolution into electrochemical formats would enable digital, objective and quantitative detection but have been plagued by the chemical and mechanical fragility of nitrocellulose membranes which do not allow printing of electrodes on the surface.^14^ Unlike visual interpretation or image-based analysis, which can be influenced by lighting and user dependent variability leading to poor analytical performance, electrochemical LFAs would convert chemical signals into measurable electrical outputs, enhancing performance and robustness in diverse settings.^15,16^

In this work, we report ELLA, a portable, scalable, and quantitative digital LFA tailored for point-of-use testing of pathogens, overcoming limitations associated with existing methods of detection (**Fig. 1**). ELLA introduces three new technologies to enable, high-performance, quantitative and digital testing with geolocation capabilities at the point-of-use: (i) Near-field communication (NFC)-enabled wireless and battery-free, fully disposable LFA cassettes with integrated electronic measurement circuitry; (ii) Metallic pin electrodes to measure the electrochemical signals quantitatively and; (iii) gold nanoparticles (AuNPs) modified to produce an electrochemical signal at the test line. The ELLA technology platform was validated both in the laboratory in the United Kingdom and under field conditions in Tanzania, East Africa and benchmarked against RT-qPCR and ELISA. We also developed a deep-learning–based image recognition model (DeepELLA), which uses smartphone photographs of leaves labeled as healthy or infected based on ELLA results. By leveraging these molecularly confirmed images, DeepELLA can be retrained directly in the field and rapidly deployed at scale for CBSD and other emerging pathogens, albeit with reduced analytical performance compared to ELLA-based molecular testing. Together these efforts show how data-driven models can complement the core assay, enhancing decision support and enabling scalable digital surveillance. Combined with its disposable, rapid, and cloud-connected design, ELLA has the potential to transform plant disease monitoring and provide critical support to government systems and smallholder farmers in vulnerable regions.

## Results and Discussion

### Modifying the classical lateral flow assay

A classical lateral flow test relies on sequential events of molecular recognition that culminate in a visible colorimetric signal. When a sample of liquid is introduced to a test device, the sample flows along a thin test strip, interacting with the recognition molecules that have been previously added to the membrane (**Fig. 2A**). Typically, antibodies are used to recognize the target molecules and are conjugated to nanoparticles to generate a visual signal, most often seen as the appearance of lines on the strip. When creating ELLA, we wanted to preserve the original LFA format as much as possible to reduce the barriers to scalability in terms of production and technological adoption. Furthermore, because ELLA had to operate reliably under field conditions (*e.g.,* variable temperatures and humidity), we avoided enzyme-based reporters and instead employed stable, non-biological labels compatible with electrochemical detection. For these reasons, we chose to modify the AuNPs used in classical LFAs with electroactive molecules to enable electrochemical transduction.

**Figure 1:**
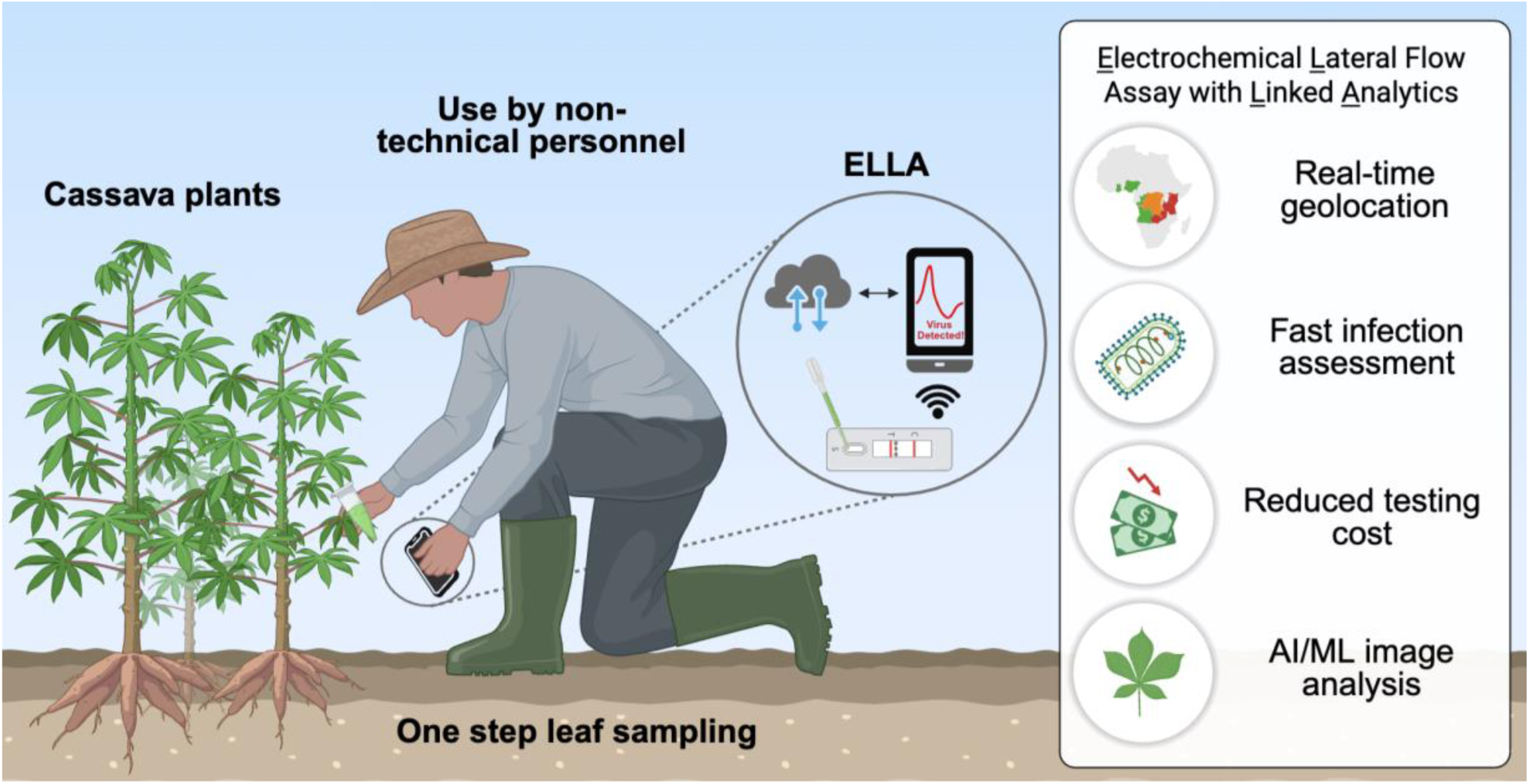
Schematic illustration of the use of the **ELLA** (**E**lectrochemical **L**ateral flow assay with **L**inked **A**nalytics), demonstrating its key benefits for field diagnostics. The device integrates an LFA platform with electrochemical sensing, allowing for precise, quantitative measurements. The diagram highlights the streamlined sample application, automated data capture, and wireless transmission via NFC to mobile devices for real-time analysis and remote monitoring. Key advantages include portability, affordability, and ease of use in resource-limited settings, offering a robust solution for early disease detection and data-driven management strategies in agricultural applications. Figure created with BioRender.com.

**Figure 2:**
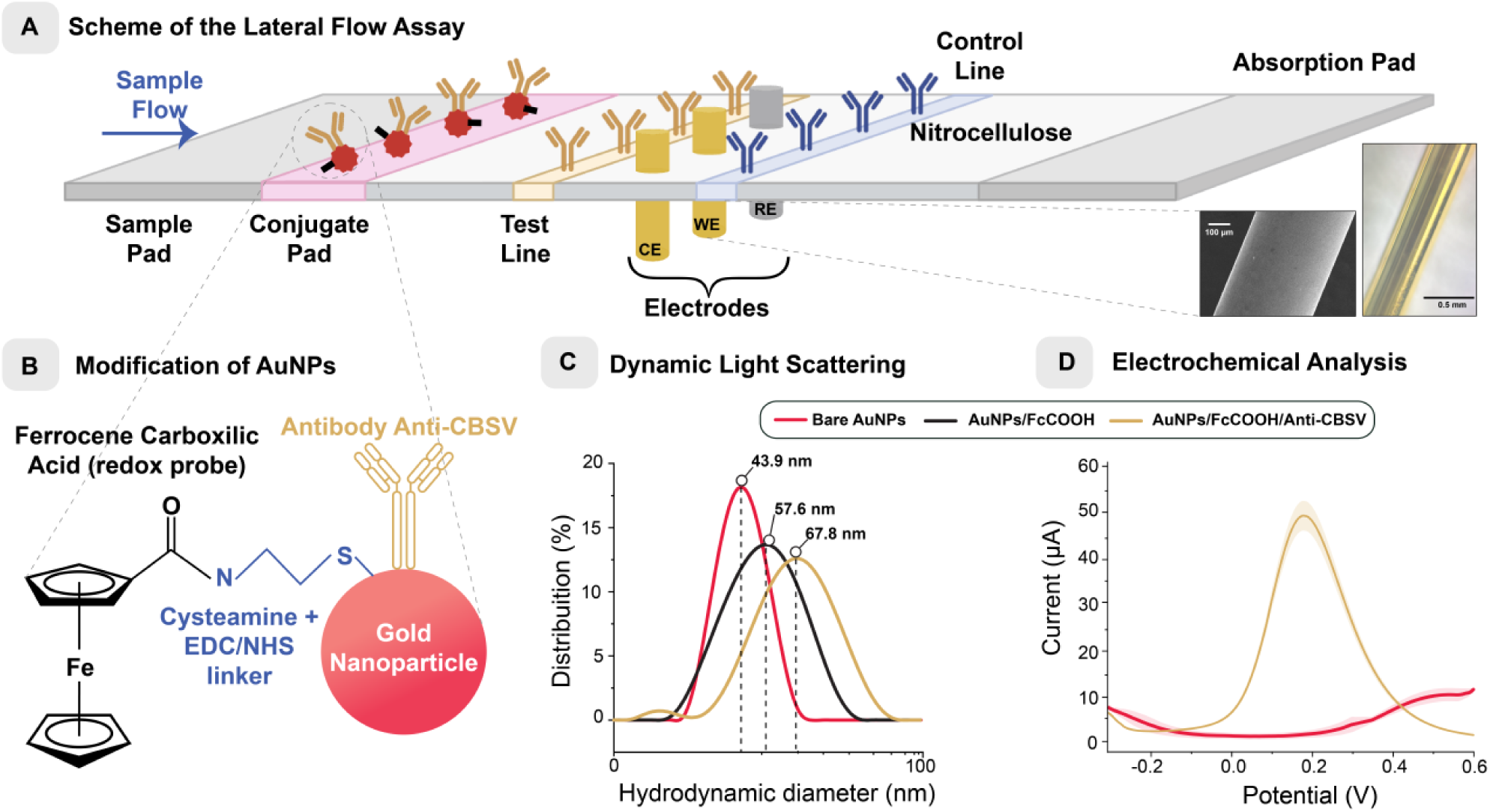
Modification of the traditional LFA: **(A)** Scheme of the LFA utilized in this work, with its typical constituents and the addition of labelled nanoparticles and electrodes. In the inset, electron micrograph and light microscopy images of the needles used as electrodes. (**B**) Scheme of the modified AuNPs designed for the detection of the CBSV, illustrating the functionalization process that enhances specificity and enables electrochemical detection (not to scale). (**C**) Dynamic Light Scattering analysis confirmed the successful modification of nanoparticles, evidenced by changes in their hydrodynamic diameter. (**D**) Square wave voltammograms showcasing the electrochemical behavior of the modified nanoparticles (n = 3).

Gold nanoparticles (⌀ = 40 nm, purchased from BBI Solutions, UK) were modified with ferrocene carboxylate before conjugation with capture antibodies against the envelope protein of the CBSV (**Fig. 2B**). Ferrocene is a low-cost, stable, near ideal reporter because the Fc/Fc⁺ redox couple undergoes a fast and reversible one-electron transfer with a well-defined mid-range potential across many solvents.^17^ We selected ferrocene carboxylate rather than unmodified ferrocene because the carboxyl group improves solubility in polar solvents and enables covalent coupling to amine-functionalized AuNPs. To achieve covalent attachment, AuNPs were functionalized with cysteamine, activated with EDC/NHS chemistry, and subsequently coupled to ferrocene carboxylate as the electroactive label. The resulting ferrocene-modified AuNPs were then conjugated, via adsorption, to anti-CBSV antibodies (obtained from the Leibniz-Institut DSMZ, Germany), which were originally developed for a sandwich ELISA and can recognize both CBSV and UCBSV. In the ELLA format, the complementary antibody from the same ELISA pair was immobilized on the test line, enabling capture of the antigen–nanoparticle complexes and generation of the electrochemical signal.

We monitored each step of the reaction process through aggregation testing and dynamic light scattering (DLS) analysis to ensure a stable colloidal suspension was formed. The aggregation test is designed to trigger the precipitation of AuNPs upon the addition of a salt solution, usually sodium chloride, causing the suspension to shift from a stable red-pink colloid to a greyish-purple hue. The AuNPs remained stable throughout all modification steps, even after the addition of a 1M sodium chloride salt solution (**Fig. S1**), demonstrating the formation of a stable colloid suitable for biological recognition assays. DLS analysis confirmed the progressive increase in the hydrodynamic diameter of the functionalized AuNPs after each step of modification (**Fig. 2C**). Bare AuNPs had an average diameter of 43.9 nm with a zeta potential of –21.6 mV, consistent with their expected colloidal stability.^18^ Following functionalization with ferrocene, the hydrodynamic diameter increased to 57.6 nm, while the zeta potential shifted to – 27.0 mV. The apparent 14 nm increase can be attributed not only to the molecular size of the ferrocene linker itself but also to the formation of a solvation layer and changes in the nanoparticle’s effective diffusion boundary after surface modification. Small-molecule grafting often increases the measured hydrodynamic diameter because DLS captures the particle plus its associated solvent and counterion shell, which becomes more structured when polar or charged ligands are introduced. ^19^ The concurrent shift to a more negative zeta potential further supports the successful attachment of ferrocene derivatives, enhancing electrostatic stabilization. Subsequent conjugation with antibodies further increased the mean hydrodynamic diameter to 67.8 nm, while the zeta potential shifted to –11.8 mV, consistent with the partial neutralization of surface charge by protein adsorption.

Next, we drop cast the colloidal suspension containing the ferrocene functionalized AuNPs onto commercial screen-printed electrodes to confirm functionalization and electroactivity through square wave voltammetry.^20^ The modified AuNPs showed a characteristic redox peak around +0.2 V ^21^ (**Fig. 2D**), indicating that the voltametric properties of ferrocene was preserved, and the conjugate solution could be used for electroanalysis.

### System architecture and operation of ELLA

The ELLA measurement platform comprises several key hardware and software components (**Fig. 3A**): i) an NFC-enabled smartphone with a custom mobile application, ii) a disposable 3D-printed test cassette housing the readout electronic printed circuit board (PCB) connected to the test strip, iii) cloud backend for online storage of the test results along with metadata concerning each measurement such as the location. The PCB features a planar loop antenna with a resonant frequency of 13.56 MHz, an NFC microchip (SIC4341 sensor interface, from Silicon Craft PLC, Thailand), and connectors for interfacing with the electrodes for performing electroanalysis (**Fig. S2 and S3**). The SIC4341 chip is a 228-byte ISO14443A passive NFC-Forum Type 2 Tag, capable of performing functions of a potentiostat and interfacing with an electrochemical sensor. SIC4341 supports both 2 and 3-electrode electrochemical measurement configurations and is compatible with amperometry and voltammetry techniques. A built-in 10-bit analog-to-digital converter (ADC) converts the electrochemical signals measured into a digital number. When an NFC-enabled smartphone is brought near the loop antenna, the radio frequency power from the magnetic field induced activates the SIC 4341 NFC microchip which also turns on an LED on the PCB to indicate operation. Once powered on wirelessly through inductive coupling, SIC4341 sensor interface begins initialization, activates peripheral devices, and enters a standby mode awaiting further operational commands from the smartphone over NFC. We designed a custom 3D-printed phone case to simplify alignment of the loop antenna in the ELLA cassettes with the loop antenna of the mobile phone. Otherwise, the operator will need to hold the phone or the cassette steady during the measurement. We also developed a custom mobile application (**Fig. S4**) to enhance user experience and data management. The application was built using Flutter (version 2.8.1) with Dart in Android Studio and is currently deployed on Android platforms, with future cross-platform extensions planned. The app features two distinct interfaces: a simplified “non-expert” mode with step-by-step guidance, and an “expert” mode offering advanced parameter customization. It supports both online and offline functionality, utilizing MongoDB Cloud Database for cloud storage and centralization, which is important for performing data analytics.

**Figure 3:**
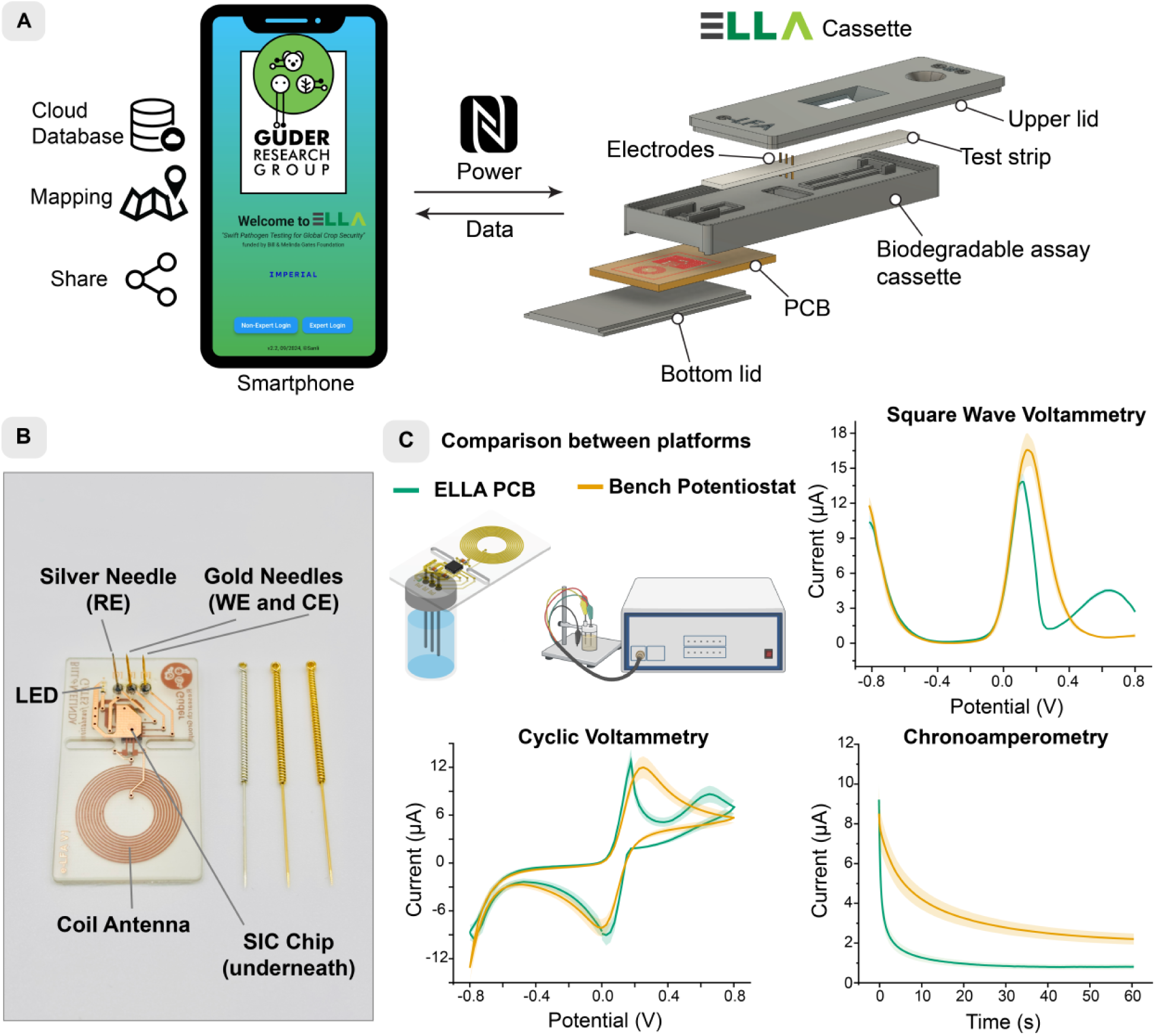
(**A**) Components of the ELLA: a custom-made smartphone app connected to the electrochemical lateral flow assay cassette via NFC technology. (**B**) Photos of the ready-to-use PCB and the needles used as electrodes. (**C**) Comparison of the developed PCB against a benchtop potentiostat (Autolab PGSTAT204, Metrohm) using standard electrochemical techniques in a ferro/ferricyanide redox probe solution (n = 3).

### Characterization of needle electrodes

Gold- and silver-plated acupuncture needles (**Fig. 3B**) were selected as electrodes for their low cost, robustness, and ease of integration into nitrocellulose membranes without damaging the substrate. Using [Ru(NH₃)₆]³⁺/²⁺ as an outer-sphere redox probe ^22^, the needle electrodes exhibited diffusion-controlled, reversible behavior comparable to commercial screen-printed electrodes, with a larger electroactive area attributed to their continuous metallic surface (**Fig. S6**). To evaluate the electronic interface, the needle electrodes were connected to the NFC-powered SIC4341 PCB potentiostat and benchmarked against a benchtop instrument using the [Fe(CN)₆]³⁻/⁴⁻ redox couple ^23^ (**Fig. 3C**). The PCB device produced slightly lower currents and displayed mildly clipped triangular CV peaks, reflecting expected limitations in compliance and bandwidth relative to the benchtop potentiostat (Autolab PGSTAT204, cost ∼US$13,000) but maintained similar voltametric profiles. An occasional secondary anodic feature was observed, attributed to capacitive and surface effects rather than a distinct redox process. Sensitivity analysis using square-wave voltammetry showed that both systems produced comparable calibration curves across a range of [Fe(CN)₆]³⁻/⁴⁻ concentrations (**Fig. S7**). The peak currents followed a logarithmic dependence on concentration with nearly identical fits (R^2^ = 0.975 for the PCB; R^2^ = 0.976 for the bench potentiostat), confirming consistent quantitative response and diffusion-limited behavior. Overall, the small single-chip potentiostat (1.2 × 1.2 mm, cost ∼US$0.20) achieved laboratory-grade electrochemical performance at over four orders of magnitude lower cost, validating both the acupuncture needles and embedded electronics as reliable, scalable components for the ELLA platform.

### Validation of the electrochemical lateral flow assay

We evaluated the feasibility of integrating the needles as detection electrodes within the LFA (**Fig. 4A**): The electrodes could monitor the capillary-driven transport of the electrochemical redox probe along the test strip and produce a reliable contact with the solution. The electroanalytical current produced by square wave voltammetry (SWV) exhibited strong time dependence, consistent with mass transport and electrolyte depletion at the electrode–liquid interface. The redox current increased substantially five minutes after the addition of the liquid sample to the device then dropped markedly potentially due to the combined effects of: (i) depletion of redox molecules near the needles, (ii) reduced conductivity of the liquid front because of flow, and (iii) diminished double-layer capacitance as the electrolyte reservoir becomes exhausted.^24^ Based on these observations, we standardized our protocol to initiate the electrochemical measurements five minutes after the application of the sample.

**Figure 4:**
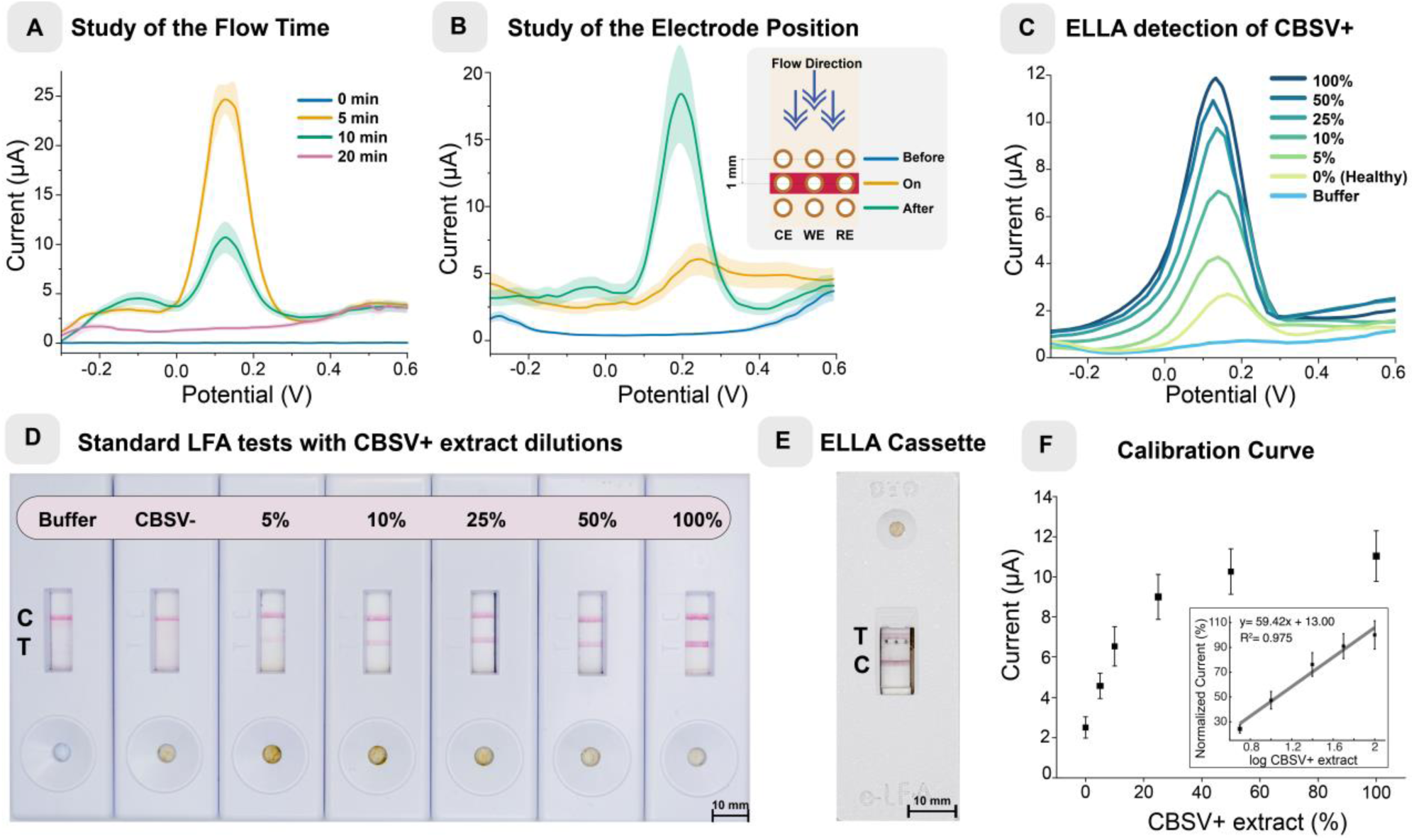
Electrochemical lateral flow test validation: (**A**) Evaluation of the flow time onto the electrochemical signal of the ELLA. (**B**) Study of the position of the test line with the electrodes. (**C**) Response of the ELLA with different dilutions of CBSV positive control. (**D**) Photograph of traditional visual LFAs with different dilutions of a positive sample. (**E**) Photograph of an ELLA cassette after testing a positive sample. (**F**) Calibration curve of the raw data obtained from the ELLA, in the insert the normalized data.

We assessed how the placement of the electrodes influenced the quality of the electrochemical signals (**Fig. 4B**). Contrary to our expectations, positioning the needle electrodes directly through the test line yielded suboptimal performance. This configuration likely introduced increased charge-transfer resistance due to the higher density of immobilized and captured proteins at the electrode–sample interface. In contrast, placing the electrodes immediately downstream from the test line produced the most robust electrochemical response, likely facilitating more efficient electron transfer between the electroactive probe retained on the accumulated AuNPs and the surface of the electrodes. We, therefore, placed all three electrodes immediately after the test line to maximize stability and reproducibility of the signals acquired in all subsequent experiments.

After characterizing each element of ELLA in isolation, we began the construction of the LFA-based test for the electrochemical detection of CBSV (**Fig. 4C**). First, we validated the biological reagents to construct the assay work as intended by creating a classical, colorimetric LFA (**Fig. 4D**). To test the assay, freeze-dried samples of CBSV-infected cassava leaves (obtained from the Leibniz Institute DSMZ, Germany) were re-suspended according to the instructions by the supplier prior to use (details in SI). Because purified standards containing the CBSV coat proteins were unavailable, calibration was performed using dilutions of the infected plant extract in healthy plant extract, with healthy plant tissue serving as the negative control and running buffer as the blank. The same samples were tested in our ELLA platform (**Fig. 4E**).

Using the redox current as the analytical signal, we produced a calibration curve relating the current intensity to the dilution of the positive sample (**Fig. 4F**). For quantitative comparison across runs, the current responses were normalized according to the following equation:

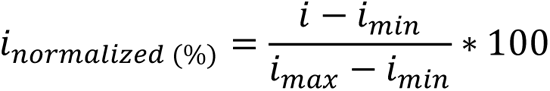

where *i* is the peak current of the SWV measurement, *i_min_* is the mean current response of the negative control and *i_max_* is the mean current response from the positive control. LFAs usually present a plateau response in high concentrations, so after applying the log function, we could linearize the response according to the equation *i_normalized_* = 59.42*log(CBSV%) + 13.00 (R^2^ = 0.975). This correlation confirms that ELLA provides a quantitative readout across a physiologically relevant concentration of infected material. The limit of detection (LOD) was 1.19% leaf extract dilution, calculated as three times the standard deviation of the negative control, which yielded a current of 4 µA. Currents exceeding this threshold were classified as positive by the ELLA software. As shown in the next section, more advanced analysis (such as the Receiver Operating Characteristics) would allow us to determine a more optimized threshold hence a lower LOD.

Because our assay detects viral coat proteins rather than viral RNA, and no purified CBSV protein standard is available, we could not express the LOD in protein concentration or genome copies as is customary for molecular assays. Instead, we determined the LOD in terms of amperometric current and relative dilutions of a well-characterized positive control. Although this approach does not allow a direct comparison with molecular tests in terms of absolute sensitivity, it aligns with the practical context in which ELLA is intended to operate. The platform was not designed to surpass RT-qPCR in analytical sensitivity, but to provide a rapid, low-cost, and field-deployable surveillance tool that complements centralized laboratory methods. Notably, ELLA outperformed ELISA (**Table S1**), detecting dilutions of the positive control below the ELISA detection limit.

As ELLA is meant to be used under field conditions without access to refrigeration, we also characterized the stability of the assay when kept at room temperature (25 °C ± 1). ELLA maintained 81% of the initial response after 150 days of storage (**Fig. S10**), indicating that the conjugated AuNPs and other elements within the assay including the electrodes, electronics, biological reagents etc., are robust enough for medium-term storage without refrigeration.

### Field testing in rural Tanzania, East Africa

ELLA is intended to be used for detecting CBSD in the field hence, we conducted a series of experiments in Chambezi (Bagamoyo, Pwani region, Tanzania) to validate the entire platform including the electrochemical molecular assay, electronics, mechanical and software components (**Fig. 5A**). The Android-based mobile application enables the operation of ELLA by less technical individuals, such as extension officers and farmers with just a few taps. Unlike ELISA and RT-qPCR, which require expert personnel for performing the test and can take up to hours or even days, running a test with ELLA from sample collection to result can be as little as 10 minutes (**Fig. 5B**). The preparation of the sample and running the test consists of three simple steps: (i) cassava leaves are sampled by pinching with the cap of the tube provided (1.5 mL microtube) to collect three leaf disks; (ii) the disks are mechanically disrupted by shaking the tube, which also contained steel beads, for one minute; and (iii) approximately 80 µL (four droplets) of the resulting extract is dispensed into the ELLA cassette. All required reagents and consumables are supplied in a single kit (**Fig. 5C**).

When using the mobile application, in addition to performing electroanalysis of the sample extracted from cassava leaves, we also captured the geolocation of the test, allowing us to monitor all field experiments in real-time from our university offices in London, UK as part of the team performed the experiments in Tanzania. We recorded every sampling location, including our field experiments in Tanzania, laboratory tests in London, and CBSD-free cassava tests in Brazil. All test results performed using the ELLA platform could be visualized through an interactive web app or Android mobile application – both developed by our group (**Fig. 5D**).

**Figure 5.**
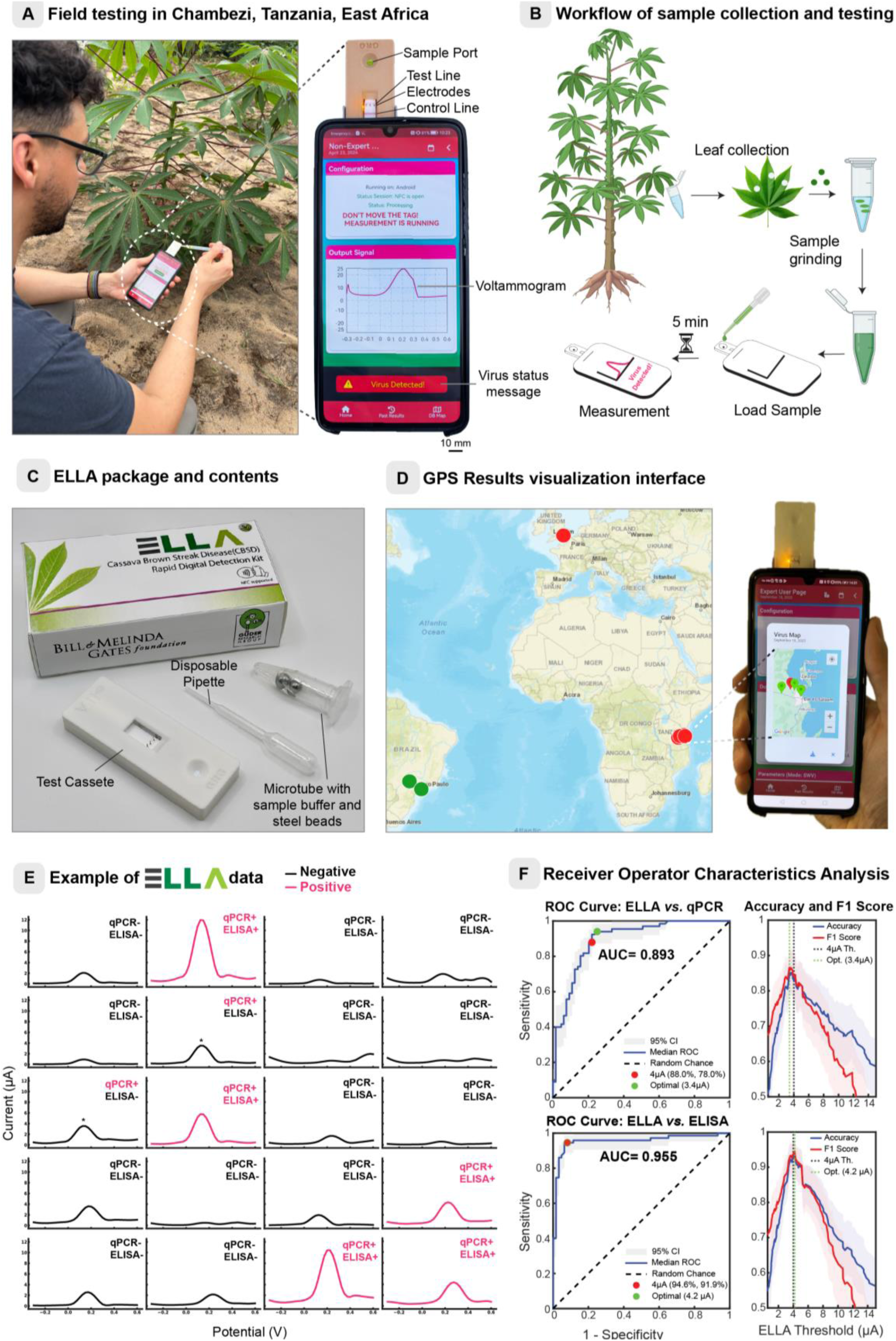
Field tests and platform validation: (**A**) Photograph of ELLA being used in a field test in Tanzania to detect CBSD in cassava plants, with a close-up of a smartphone with the custom-made Android app, connected to an ELLA cassette showing a positive test. (**B**) Sampling method for fresh cassava leaves: fresh leaves are collected and placed into microtubes, where they are thoroughly crushed with a buffer solution with steel beads. Following this, approximately 80 µL of the resulting solution is applied to the ELLA cassette for testing. After 5 minutes the results are displayed. (**C**) Photograph of an ELLA package and its contents: a test cassette, disposable pipette and a microtube containing the sample buffer with steel beads. (**D**) World map showing all the locations of sample analysis, in the inset a photo of the ELLA app with the built-in map showing sampling in the Dar es Salaam and Pwani regions. (**E**) Examples of recorded voltammograms for 20 different samples alongside RT-qPCR and ELISA results. The samples marked with a * are examples of false negatives that could be detected changing the current threshold. (**F**) Receiver Operating Characteristic (ROC) curve and threshold-dependent accuracy/F1 score of ELLA compared with PCR and ELISA, showing sensitivity–specificity trade-offs and optimal cut-off points (95% CI shown).

To evaluate the performance of ELLA in the field (**Fig 5E**), we analyzed the same leaves using both ELISA and RT-qPCR back in our laboratories in Dar es Salaam (**Table S1**). Using RT-qPCR as the benchmark, we generated a Receiver Operating Characteristic (ROC) curve yielding a median area under the curve (AUC) of 0.893 (95% CI) (**Fig. 5F**). This analysis also quantified threshold-dependent variations in accuracy and F1 score, enabling us to explore sensitivity–specificity trade-offs and to mark the 95% bootstrap confidence intervals. When benchmarked against ELISA, the AUC increased to 0.955, reflecting a strong concordance between the two immunoassays. This higher correlation is expected because both ELLA and ELISA detect viral proteins, whereas RT-qPCR detects viral genetic material. Based on this analysis, the optimal detection threshold was 3.4 µA when using RT-qPCR as the reference and 4.2 µA when benchmarked against ELISA. Initially, we set the threshold at 4 µA, calculated as three times the standard deviation of the negative sample. However, these results show that the detection threshold can be refined through further sampling and analysis, and the app can be updated accordingly to improve classification performance. ELLA also demonstrated selectivity, as several collected plants infected with Cassava mosaic disease (CMD) produced no cross-reactive signal (see **Table S1**).

We also explored whether supervised machine learning could extract additional predictive value from the electrochemical measurements. A total of 125 SWV signals were processed to derive a range of features, including peak characteristics (e.g., maximum/minimum current, potential, prominence, width), statistical descriptors (mean, standard deviation, skewness), area under the curve, slope parameters, frequency-domain features from fast Fourier transform (FFT), and coefficients from polynomial fitting. Feature correlation analysis revealed 13 independent contributors, of which two were selected as the most informative for classification: maximum current and minimum current (**Fig. S11**). These features were subsequently used to train and validate multiple algorithms, including logistic regression, random forest, XGBoost, support vector machine, and multilayer perceptron. We incorporated extensive parameter tuning and cross-validation to improve learning performance, and the best-performing models reached an accuracy of 84%, F1 score of 86%, and an ROC AUC value of 90% on the test set (20% of the dataset). These results, although promising, did not exceed the performance achieved by simple thresholding of the peak current. This is likely because the two most informative features are related to current amplitude indicating that classical approaches remain effective for this dataset.

### DeepELLA: A deep learning model trained with ELLA for image-based detection of CBSD

Although clearly it is always best to perform molecular testing to confirm the presence of CBSD using ELLA, as a physical test, there may be supply related issues and other practical issues that reduce efficiency of surveillance. Furthermore, in some circumstances, the increased diagnostic performance of molecular testing can be traded off for the cost and speed of image-based analysis. We have also developed a workflow for AI-based diagnosis of infections of CBSD (**Fig. 6A**) to supplement extension officers that solely rely on visual inspections for diagnosis. ELLA-based molecular testing can be performed in the field rapidly and the results of these tests, along with the images of leaves from the plants, can be used to train a model that can run on a smartphone. In other words, ELLA can be used to generate ground truth data for training the model. To validate this approach, we imaged 109 of the leaves used for the molecular tests with a smartphone, comprising 53 healthy and 56 diseased samples (**Fig. 6B**).

Because we want this approach to work with a small number of images to reduce the number of molecular tests, we used a transfer learning method. For this, we opted to use ResNet-18, a pre-trained convolutional deep learning network consisting of 18 layers and modified the final layer by training with 87 images (80%). We evaluated the performance of the model using 22 test images (20%) that were not included in the training dataset. The model (DeepELLA) had an accuracy of 0.86, precision of 0.83 and an F1 score of 0.86 (**Fig. 6C**) identifying the plants with CBSD. The model is a substantial improvement to visual inspections with the naked eye such that two of the leaf samples (as shown in Fig. 6C), that were correctly classified by DeepELLA, appear practically the same. Especially, less experienced field workers would struggle to correctly identify the infected plant.

**Figure 6.**
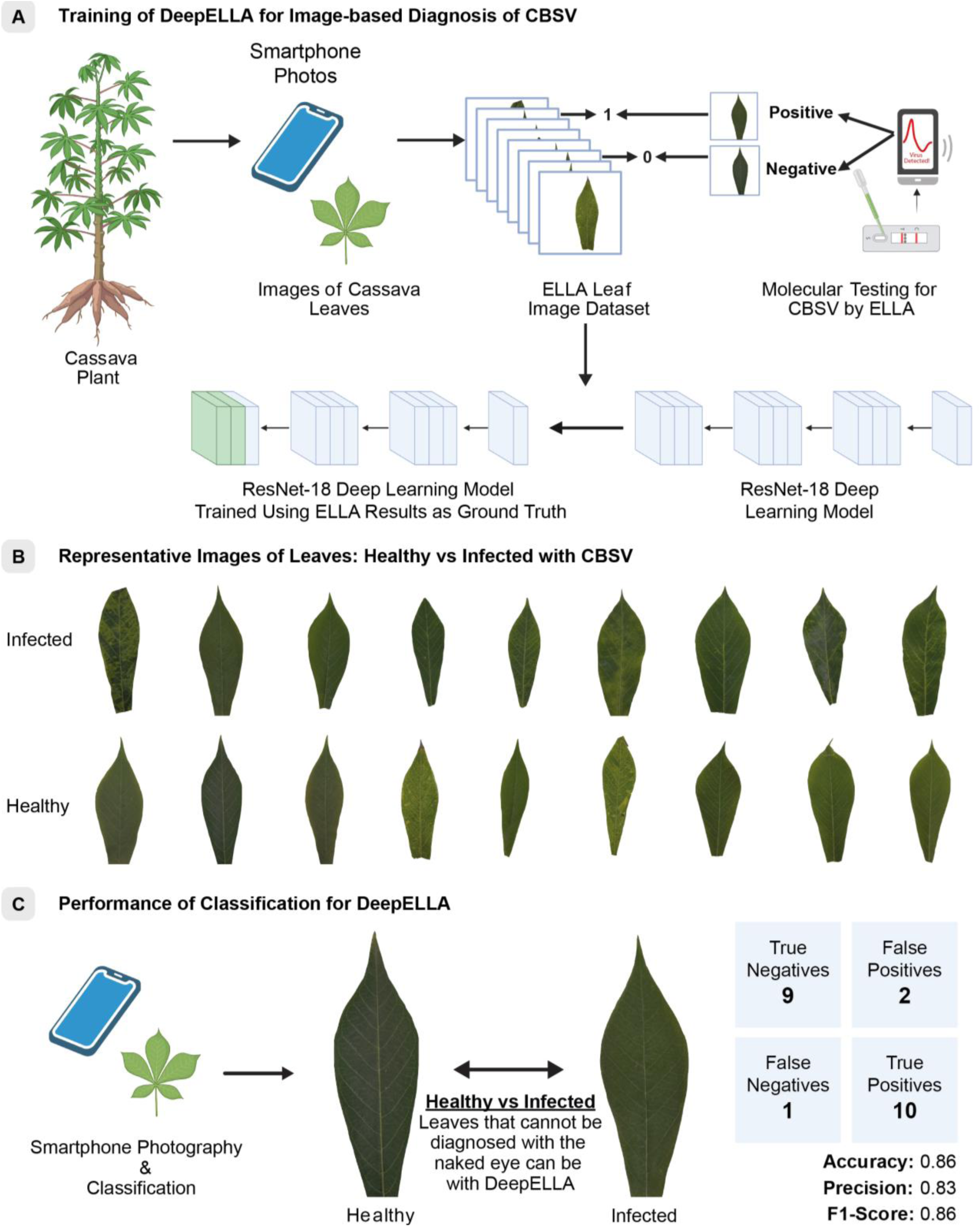
Using the deep learning method for ELLA prediction (DeepELLA). (**A**) General Scheme of the DeepELLA. (**B**) Representative photos of cassava leaves used for the analysis. (**C**) Performance of DeepELLA

## Conclusions

The ELLA platform brings fully integrated, sample-to-result, digital diagnostics to the field to aid farmers, agronomists, national and international organizations to combat outbreaks of infectious diseases more effectively. ELLA complements the established laboratory-based molecular diagnostics, performing similar or better than ELISA in terms of analytical performance but requires no additional instrumentation than a smartphone. Despite being an advanced, wireless electroanalytical technology for the molecular detection of pathogens, ELLA cassettes are low-cost and easy-to-use. A single cassette of ELLA, which contains all the biological reagents, electronics and mechanical components, costs less than US$1 for medium to high volumes. By using the results of ELLA to train an image recognition model, the effective cost of a single test can be traded off for lower analytical performance and speed of deployment at scale, especially important for novel pathogens. Furthermore, model training can be expanded to drone imaging for even larger scale surveillance – though we anticipate lower analytical performance compared to ELLA or smartphone-based image analysis.

Despite the advances we made in constructing a portable and connected analytical platform validated in the field in resource-limited settings, the ELLA cassettes have at least three disadvantages: (i) when stored at room temperature, the assay maintained most of its performance but the electroanalytical current decreased by 19% over 150 days. However, the level of stability is still sufficient when compared with typical antibody-based assays.^25^ In its present form, the assay could be stored and distributed in resource-limited settings without a strict cold chain. More optimized storage buffers, smart packaging, and biorecognition elements can, however, extend shelf life and ensure consistent performance over longer periods; (ii) the assay developed for the detection of CBSD relies on antibodies, negatively impacting affordability and access. Antibodies, however, can be potentially replaced with other biorecognition elements such as aptamers, which are more affordable and scalable as they are produced synthetically; (iii) although ELLA cassettes are battery-free, each cassette consists of a biodegradable housing made of PLA but also small amount of electronic materials which may require more specialized recycling or disposal.

In this work, we tailored the ELLA platform for the detection of CBSD due to the severity of its impact in East Africa. Nevertheless, the system is inherently modular and versatile: by simply changing the biorecognition molecules, the same platform can be adapted to detect other pathogens and diseases. Moreover, the use of electrochemical transduction provides robust quantitative readouts, further enhancing the analytical utility of the platform beyond traditional lateral flow assays. The assay can be applied to different sample mediums from blood and drinking water to food matrices for applications in healthcare, environmental monitoring and food safety. In the future, the ELLA platform can be a powerful tool for capturing a large range of biological and chemical data from complex systems to produce unique datasets. By leveraging artificial intelligence, we can get so much more from a single measurement to understand the underlying dynamics, relationships and trends between seemingly unrelated phenomena, allowing us to understand our world at a fundamentally new level.

## Methodology

All reagents were obtained from commercial suppliers (primarily Sigma–Aldrich/Merck, UK) and used without further purification unless otherwise noted.

### Gold nanoparticles modification

We started with 1 mL of optical density (OD) 5 citrate-stabilized AuNPs (40 nm diameter, BBI Solutions), which we centrifuged at 5000 × g for 10 minutes and resuspended in 2 mL of 10 mM HEPES buffer (pH 7.4) containing 0.01% (v/v) Tween 20. We then added 50 µL of 10 mM cysteamine progressively into the nanoparticle solution and incubated the mixture under agitation for 15 minutes. After incubation, we centrifuged the solution again at 5000 × g for 10 minutes, discarded the supernatant, and resuspended the pellet in 2 mL of the same HEPES buffer. Next, we conjugated the amine-functionalized AuNPs with ferrocene carboxylate (FcCOOH) using carbodiimide chemistry. We first dissolved FcCOOH as a 5 mM stock in anhydrous ethanol and diluted it in 10 mM MES buffer (pH 5.5) to a final concentration of 1 mM. We then added freshly prepared 1-ethyl-3-(3-dimethylaminopropyl)carbodiimide (EDC, 10 mM final) and sulfo-N-hydroxysuccinimide (sulfo-NHS, 20 mM final) to the FcCOOH/MES mixture and allowed the reaction to proceed for 15 minutes at room temperature to generate the NHS ester. We added the activated FcCOOH solution dropwise to the AuNP suspension to reach a final FcCOOH concentration of 200 μM and incubated the mixture under gentle agitation for 60 minutes at room temperature to enable amide bond formation with the nanoparticle surface amines. To quench residual esters, we added ethanolamine (5 mM, pH 8.0) and incubated for 15 minutes. Finally, we purified the conjugated AuNPs by centrifugation (5000 × g, 10 minutes), discarded the supernatant, and resuspended the pellet in 10 mM HEPES (pH 7.4) containing 0.01% (v/v) Tween-20. We obtained anti-CBSV antibodies from the Leibniz-Institut DSMZ (Braunschweig, Germany) and conjugated them to the ferrocene-modified AuNPs by adsorption. We mixed the AuNPs with the anti-CBSV antibody solution (DSMZ AS-1153) diluted in MOPS buffer (pH 8.0) to a final antibody concentration of 5 μg/mL, then incubated the solution for 10 minutes at room temperature with gentle inversion. To block non-specific binding sites, we added bovine serum albumin (BSA) to a final concentration of 20 μg/mL and incubated for another 30 minutes. We centrifuged the mixture at 5000 × g for 10 minutes, carefully removed the supernatant, and resuspended the pellet in gold drying buffer (0.1 M Tris, 3% BSA, 5% sucrose, 1% Tween 20, pH 9.4) to a final OD of 10. We evaluated each modification step using a Zeta Sizer spectrophotometer and a gold aggregation test. To verify electrochemical activity, we drop-cast 10 μL of the colloid solutions onto screen-printed gold electrodes (Metrohm DropSens 220AT) and performed square wave voltammetry (–0.3 V to +0.6 V, amplitude 25 mV, frequency 5 Hz) on a Metrohm Autolab PGSTAT204 potentiostat using PBS as the electrolyte. In a separate step, we prepared the control line conjugate. We centrifuged 1 mL of goat anti-biotin AuNPs (OD 10, BBI Solutions) at 5000 × g for 10 minutes, removed the supernatant, and resuspended the pellet in the previously prepared AuNP–anti-CBSV conjugate solution. We then deposited this nanoparticle conjugate onto a glass fiber membrane (300 mm × 22 mm, Millipore) using the AirJet pump of a BioDot dispenser at a flow rate of 0.5 μL/mm

### Design of the receiving antenna and user interface

We designed a custom planar antenna with a 15 mm diameter and 10 turns on a transparent epoxy substrate to enable wireless power transfer through the NFC reader. The antenna had an inductance of 1.32 µH and a quality factor exceeding 50 at the 13.56 MHz operating frequency (Fig. S3). We calculated the required external tuning capacitor as 54 pF to complement the 50 pF internal capacitor of the SIC4341 chip. This chip integrates a potentiostat for current measurements, which we controlled through our custom mobile app (Fig. S4). We developed the app with separate user interfaces for “expert” and “non-expert” users, allowing flexible measurements and streamlined data sharing via MongoDB Atlas, thereby enhancing the usability and field diagnostic capabilities of the ELLA device.

### Electrochemical tests

We performed electrochemical measurements using a three-electrode configuration, repurposing acupuncture needles as miniaturized electrodes due to their favorable size, conductive coating, and mechanical robustness. We used gold-plated stainless-steel needles (CD7) as the working and counter electrodes, and a silver-plated stainless-steel needle (CD20) as the reference electrode, all sourced from Wujiang City Cloud & Dragon Medical Device Company, China. Each needle had a diameter of 0.40 mm and was used without further surface modification. We soldered the electrodes directly onto our custom-designed PCBs hosting the SIC4341 single-chip potentiostat (Silicon Craft Technology), which enabled fully integrated, miniaturized electrochemical control and measurement. To benchmark our system against a standard benchtop potentiostat, we submerged 5 mm of the needle electrodes in a ferri/ferrocyanide redox probe (2 mM in PBS) and performed three electrochemical techniques: square wave voltammetry (–0.8 V to +0.8 V, amplitude 25 mV, frequency 5 Hz), cyclic voltammetry (–0.8 V to +0.8 V, 50 mV/s), and chronoamperometry (+0.2 V, 60 s). We recorded the data using both our custom-made smartphone app (Huawei P30) and the NOVA software on a Metrohm Autolab PGSTAT204 potentiostat. To assess the feasibility of using the needles as detection electrodes within the LFA, we constructed a blank assay (sample, conjugate, and adsorption pads plus the nitrocellulose without biomolecules). We applied 80 μL of ferri/ferrocyanide redox probe solution (5 mM in PBS with 0.1% Tween 20) to the sample pad and measured the electrochemical signal on our smartphone app using square wave voltammetry (–0.3 V to +0.6 V, amplitude 25 mV, frequency 5 Hz) at different time intervals (0, 5, 10, and 20 minutes after application). To determine the optimal electrode placement, we positioned the PCB needle electrodes at three locations: directly under the test line, 1 mm downstream, and 1 mm upstream. After adding 80 μL of electrolyte solution (PBS with 0.1% Tween 20) and waiting five minutes, we performed square wave voltammetry using our app (–0.3 V to +0.6 V, amplitude 25 mV, frequency 5 Hz).

### Lateral Flow Assay Construction

Fig. S8 shows the workflow of the LFA manufacture. For the control line, we prepared a 1 mg/mL solution of biotinylated BSA in PBS containing 2% sucrose. For the test line, we prepared a 1.5 mg/mL solution of anti-CBSV antibody (DSMZ AS-1153/1) in PBS containing 2% sucrose and 10% ethanol. We applied both solutions onto nitrocellulose membranes (FF80HP Plus, Cytiva) with a BioDot plotter at a rate of 0.2 μL/mm, positioning the control line at 13 mm and the test line at 7 mm from the membrane edge. We then dried the membranes at 40 °C for 30 minutes. Next, we assembled the lateral flow test onto a backing paper (300 mm × 60 mm, Cytiva). We glued the 20 mm-wide nitrocellulose membrane first, followed by the conjugate pad (the previously conjugated-printed glass fiber membrane, 300 mm × 20 mm, Millipore), and finally the adsorption pad at the end of the assay (Whatman CF7, 22 mm wide, Cytiva). We sliced the assembled cards into 6 mm-wide strips, stored them with silica gel pouches, and allowed them to mature overnight at 37 °C. The following day, we aligned the PCB needles with the test line using a custom mold (**Fig. S9**), pinched them into the back of the strips, and assembled the system into a 3D-printed cassette (Bambu Lab Mini A1, polylactic acid, Verbatim).

### Sample preparation and detection

For the calibration curve, we used positive and negative controls for CBSD obtained from the Leibniz Institute DSMZ (Germany). The controls consisted of freeze-dried cassava leaves, which we resuspended in extraction buffer (0.2 M Tris, 1 M NaCl, 2% PVP, and 0.1% Tween 20). For fresh samples, we cut cassava leaf disks (⌀ = 4 mm) using a microtube cap and mechanically ground them in extraction buffer (1:10 ratio of leaf weight to buffer) with steel beads for 1 minute. For both fresh and freeze-dried samples, we applied 80 μL of the resulting extract to the sample port of the ELLA. After 5 minutes, we recorded a square wave voltammogram using our smartphone app (–0.3 V to +0.6 V, amplitude 25 mV, frequency 5 Hz) on a Huawei P30. We used the peak current as the detection parameter. In parallel, we performed ELISA and RT-qPCR on all leaf samples to detect CBSV/UCBSV, following the detailed protocols reported in the Supplementary Information.

### Deep Learning Architecture

In this study, we used the Residual Neural Network (ResNet-18) architecture to predict the ELLA class from cassava leaf images. ResNet-18 is a convolutional neural network (CNN) with 18 layers, including convolutional, pooling, and fully connected layers. ^26^ We chose this architecture because it provides a good balance between model complexity and computational efficiency, making it suitable for relatively small datasets such as our collection of leaf images. For training, we resized all images to 224 × 224 pixels to match the model’s input dimensions. To reduce overfitting and improve generalization, we applied data augmentation techniques including rotation and reflection. We trained the model using the Adam optimization algorithm with an initial learning rate of 0.0001, 50 epochs, and a batch size of 32. We evaluated classification performance using accuracy, precision, and F1-score metrics.

## Acknowledgements

The authors acknowledge the Gates Foundation (Grand Challenges Explorations scheme under grant numbers OPP1212574 and INV-038695) for financial support. We are especially grateful to Jim Lorenzen and Neil Hausmann for their invaluable guidance and continued encouragement during this project. F.G. and J.M.R.F. acknowledge Research England International Science Partnerships Fund (Reference: G08259) and the Biotechnology and Biological Sciences Research Council Impact Acceleration Account (Reference: PSR417). F.G. and A.S. acknowledge Research England International Science Partnerships Fund (Reference: G08288) and Innovate UK BMC (Reference: PA2131). F.G. and L.G.-M. thank the European Union’s Horizon 2020 research and innovation program under the Marie Sklodowska-Curie grant agreement No 101025390. F.G. and S.O. acknowledge Bezos Earth Fund through the Bezos Centre for Sustainable Protein (BCSP/IC/001). A.G. would like to thank the TUBITAK (The Scientific and Technological Research Council of Türkiye), for their support under grant number: 1059B192400999. We also thank Professor Susan Seal, Gonçalo Ramalho e Silva, and Sophie Bouvaine from the University of Greenwich for kindly providing some of the cassava samples used in this study.

## Data availability

Data will be made available on request.

## References

1. Patil, B. L., Legg, J. P., Kanju, E. & Fauquet, C. M. Cassava brown streak disease: a threat to food security in Africa. J. Gen. Virol. 96, 956–968 (2015).

2. Parmar, A., Sturm, B. & Hensel, O. Crops that feed the world: Production and improvement of cassava for food, feed, and industrial uses. Food Secur. 9, 907–927 (2017).

3. Lister, R. M. Mechanical transmission of cassava brown streak virus. Nature 183, 1588– 1589 (1959).

4. Tomlinson, K. R., Bailey, A. M., Alicai, T., Seal, S. & Foster, G. D. Cassava brown streak disease: historical timeline, current knowledge and future prospects. Mol. Plant Pathol. 19, 1282–1294 (2018).

5. Ferris, A. C., Stutt, R. O. J. H., Godding, D. & Gilligan, C. A. Computational models to improve surveillance for cassava brown streak disease and minimize yield loss. PLoS Comput. Biol. 16, e1007823 (2020).

6. Legg, J. et al. Community phytosanitation to manage cassava brown streak disease. Virus Res. 241, 236–253 (2017).

7. Carlson, C. J. et al. Pathogens and planetary change. Nat. Rev. Biodivers. 1, 32–49 (2025).

8. Robson, F., Hird, D. L. & Boa, E. Cassava brown streak: A deadly virus on the move. Plant Path. 73, 221–241 (2024).

9. Munguti, F. M. et al. Real-time reverse transcription recombinase polymerase amplification (RT-RPA) assay for detection of cassava brown streak viruses. Sci. Rep. 14, 12438 (2024).

10. Abarshi, M. M. et al. Optimization of diagnostic RT-PCR protocols and sampling procedures for the reliable and cost-effective detection of Cassava brown streak virus. J. Virol. Methods 163, 353–359 (2010).

11. Shirima, R. et al. Absolute quantification of cassava brown streak virus mRNA by real-time qPCR. J. Virol. Methods 245, 5–13 (2017).

12. Sseruwagi, P., Tairo, F. & Ndunguru, J. The Cassava Diagnostics Project (CDP): A review of 10 years of research 2008–2018. Tanzania Agricultural Research Institute (TARI)–Mikocheni, Dar es Salaam, Tanzania (2018).

13. Pedreira-Rincón, J. et al. A comprehensive review of competitive lateral flow assays over the past decade. Lab Chip 25, 2578–2608 (2025).

14. Gonzalez-Macia, L. et al. NFC-enabled potentiostat and nitrocellulose-based metal electrodes for electrochemical lateral flow assays. Biosens. Bioelectron. 251, 116124 (2024).

15. Ying, X. et al. Electrochemical Lateral Flow Immunoassay with Built-In Electrodes for Ultrasensitive and Wireless Detection of Inflammatory Biomarkers. Anal. Chem. 96, 10630–10638 (2024).

16. Parolo, C. et al. Tutorial: design and fabrication of nanoparticle-based lateral-flow immunoassays. Nat. Protoc. 15, 3788–3816 (2020).

17. Gagne, R. R., Koval, C. A. & Lisensky, G. C. Ferrocene as an Internal Standard for Electrochemical Measurements. Inorg. Chem. 19, 2854–2855 (1980).

18. Wei, X., Alam, A. R., Mo, Q. & Hernandez, R. Structure and Zeta Potential of Gold Nanoparticles with Coronas of Varying Size and Composition. J. Phys. Chem. C 129, 4204–4214 (2025).

19. Stetefeld, J., McKenna, S. A. & Patel, T. R. Dynamic light scattering: a practical guide and applications in biomedical sciences. Biophys. Rev. 8, 409–427 (2016).

20. Ribeiro, J. A., Silva, E., Moreira, P. S. & Pereira, C. M. Electrochemical Characterization of Redox Probes at Gold Screen-Printed Electrodes: Efforts towards Signal Stability. ChemistrySelect 5, 5041–5048 (2020).

21. Sabahat, S., Janjua, N. K., Akhter, Z. & Hassan, M. U. Ferrocene-functionalized gold nanoparticles: study of a simple synthesis method and their electrochemical behavior. Chem. Papers 73, 943–951 (2019).

22. Schrattenecker, J. D. et al. Hexaammineruthenium (II)/(III) as alternative redox-probe to Hexacyanoferrat (II)/(III) for stable impedimetric biosensing with gold electrodes. Biosens. Bioelectron. 127, 25–30 (2019).

23. Cassidy, J. F., de Carvalho, R. C. & Betts, A. J. Use of Inner/Outer Sphere Terminology in Electrochemistry—A Hexacyanoferrate II/III Case Study. Electrochem 4, 313–349 (2023).

24. Le Brun, G. et al. Studying ion transport dynamics in electrochemical measurements of lateral flow assays. J. Electroanal. Chem. 966, 118399 (2024).

25. Ma, H., Ó’Fágáin, C. & O’Kennedy, R. Antibody stability: A key to performance - Analysis, influences and improvement. Biochimie 177, 213–225 (2020).

26. He, K., Zhang, X., Ren, S. & Sun, J. Deep residual learning for image recognition. Proc. IEEE Comput. Soc. Conf. Comput. Vis. Pattern Recognit., 770–778 (2016).

