## Supplementary Information for "Electrochemical Lateral Flow Assay with Linked Analytics for the Surveillance of Cassava Brown Streak Disease in East Africa"

**AuNPs aggregation test.** In each step of the AuNPs modification, a 200 µL aliquot was collected and placed in a 96-well plate. The aggregation of the nanoparticles was induced by applying 40 µL of a 1M solution of NaCl. The absorbance in the gold aggregation protocol was measured at 550 nm and 600 nm to assess the levels of aggregation in the wells. The higher the A550nm/A600nm rate, the more stable the colloid. As shown in Fig. S1 the colloid was more stable after the modification with the antibody, with no visible precipitation after the addition of salt.

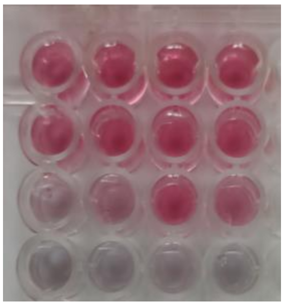

| **Sample** | **A550nm/A600nm** | | | |
|---|---|---|---|---|
| AuNPs + Water | 3.87 | 3.52 | 3.59 | 3.56 |
| AuNPs/FcCOOH/AntiCBSV + NaCl 1M | 2.62 | 3.25 | 3.53 | 3.39 |
| AuNPs/FcCOOH + NaCl 1M | 1.07 | 1.37 | 2.08 | 1.77 |
| AuNPs + NaCl 1M | 0.93 | 0.91 | 0.95 | 0.91 |

**Figure S1**: Photography of a 96 well plate with gold nanoparticles colloid modified with different molecules before (first row) and after ($2^{nd}$ to $4^{th}$ rows) the addition of 1M of NaCl (n = 4).

**Hardware Design.** We calculated the theoretical inductance value of a one-layer circular inductor using the Texas Instruments custom coil designer tool[1]. With an outer diameter of inductor ($D_{out}$) of 15 mm, a trace width (w) of 0.25 mm, a distance between traces (s) of 0.15 mm, 10 turns per layer (n) and a copper thickness of 1 oz-cu (~0.0347 mm). The resulting theoretical inductance value is 1.322 µH. This single-layer circular coil design achieves a quality factor greater than 50. As the resonance frequency for the near-field communication (NFC) tag is 13.56 MHz, we can easily calculate the external capacitor using the following formula:

$$f_{res} = \frac{1}{2\pi\sqrt{LC_{tun}}}$$

where $f_{res}$ is the resonance frequency, L is the inductance of the antenna, and $C_{tun}$ is the tuning capacitor, which is calculated as 104.205 pF. Since the internal capacitor of the SIC4341 is 50 pF, the required external capacitor would be 54.205 pF ($C_{ext} = C_{tun} - C_{int}$). We selected a 56 pF external capacitor, as it was the closest available option on the market.

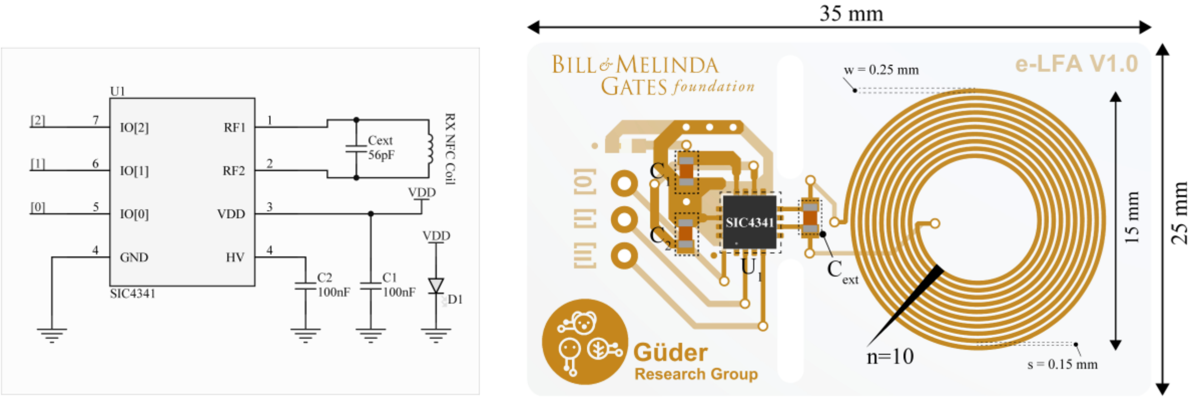


**Figure S2**: Schematic (left) and 3D PCB design (right) illustrating the receiver coil configuration. The schematic presents the functional components and connectivity of the receiver coil, while the 3D design highlights key design properties including dimensions, coil orientation, and layout on the PCB.

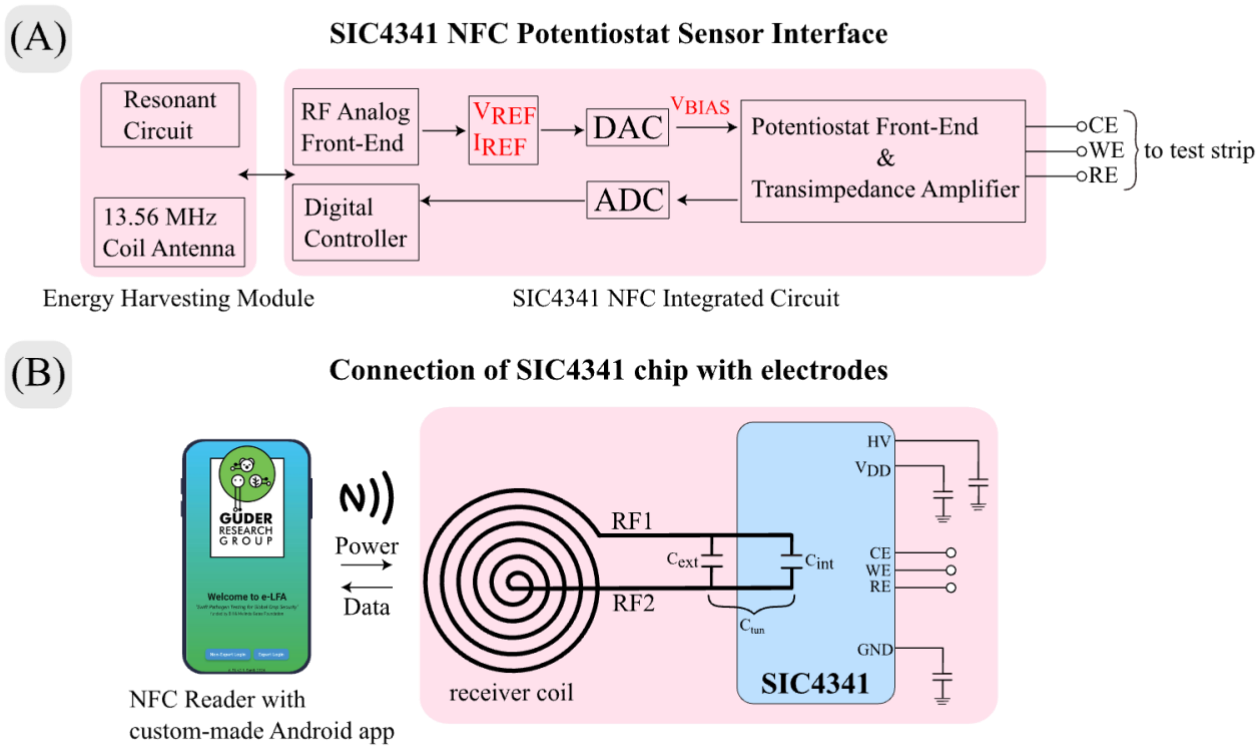


**Figure S3**: (A) The SIC4341 NFC potentiostat sensor interface, highlighting the data flow and interface setup. This figure illustrates how the SIC4341 chip, equipped with DACs, ADC, and RF AFE, communicates with connected electrodes, showing critical signal pathways and connection points for efficient data transmission. (B) Demonstrates the NFC wireless communication between a smartphone and the SIC4341 chip connected to the electrodes, facilitating seamless data transfer and power supply through NFC technology.

The SIC4341 NFC microchip incorporates an internal potentiostat designed to measure currents from chemical sensors, with a maximum input range of ± 20 μA. This chip can generate a biasing voltage between -800 mV and 800 mV for the working and reference electrodes, using dual digital-to-analog converters (DACs) with a 1.28 V full-scale voltage and 5 mV resolution. Our custom-made mobile application controls the biasing voltage of the sensor by sending commands to adjust the DAC output. The sensor current is processed by a 10-bit analog-to-digital converter and a digital signal processor and then stored in internal memory for transmission back to the smartphone.

To enhance user experience and provide visual feedback, we integrated an LED indicator into the PCB that illuminates when a stable NFC connection is established between the smartphone and the ELLA device. This feature improves usability and aids in troubleshooting. We printed the ELLA test cassettes using translucent Polylactic acid (PLA) filament, allowing users to observe the LED and receive real-time feedback on NFC connectivity while maintaining our commitment to sustainability. To ensure measurement consistency, we also designed a specialized 3D-printed phone case (demonstrated with a Huawei P30 Lite) that optimizes ELLA cassette placement and maintains uniform power distribution. These enhancements collectively improve the functionality and user experience of the ELLA device, providing clear visual cues for successful communication and facilitating proper device operation.

**Software Development.** We developed a user-friendly custom-made app interface, where we have meticulously designed two login pages tailored for "non-expert" and "expert users" (see Fig.S3). The non-expert page prioritizes simplicity, featuring a detailed instructional guide to assist users in conducting precise measurements. Upon insertion of the test cassette and initiation of the measurement process via the START button, users are prompted to input basic details such as username, file name, and type of virus (i.e., CBSV, BBTV, etc.), after which the app autonomously progresses through calibration, conversion, and result retrieval. The "expert page" on the other hand enables the user more flexibility in parameter customization, allowing selection between voltammetric (i.e. square wave voltammetry, cyclic voltammetry) or amperometric (chronoamperometry) measurements. Notably, ELLA operates seamlessly both online and offline, leveraging MongoDB Atlas cloud Database for online functionalities. This integration enables users to monitor, and share collected data, alongside a Database Map illustrating virus spread over time. Additionally, all measurements are catalogued in a calendar format for easy reference.

The cloud-based storage of measurement data enhances the functionality of the mobile app by allowing users to share results through various communication channels. Users can easily distribute their findings via email, SMS, or other messaging platforms, facilitating rapid information dissemination among researchers, healthcare professionals, or relevant stakeholders. This feature promotes collaboration and enables quick response to potential outbreaks or emerging trends in virus detection. The combination of user-friendly interfaces, flexible measurement options, and robust data-sharing capabilities makes ELLA a comprehensive tool for both field diagnostics and advanced laboratory analysis.

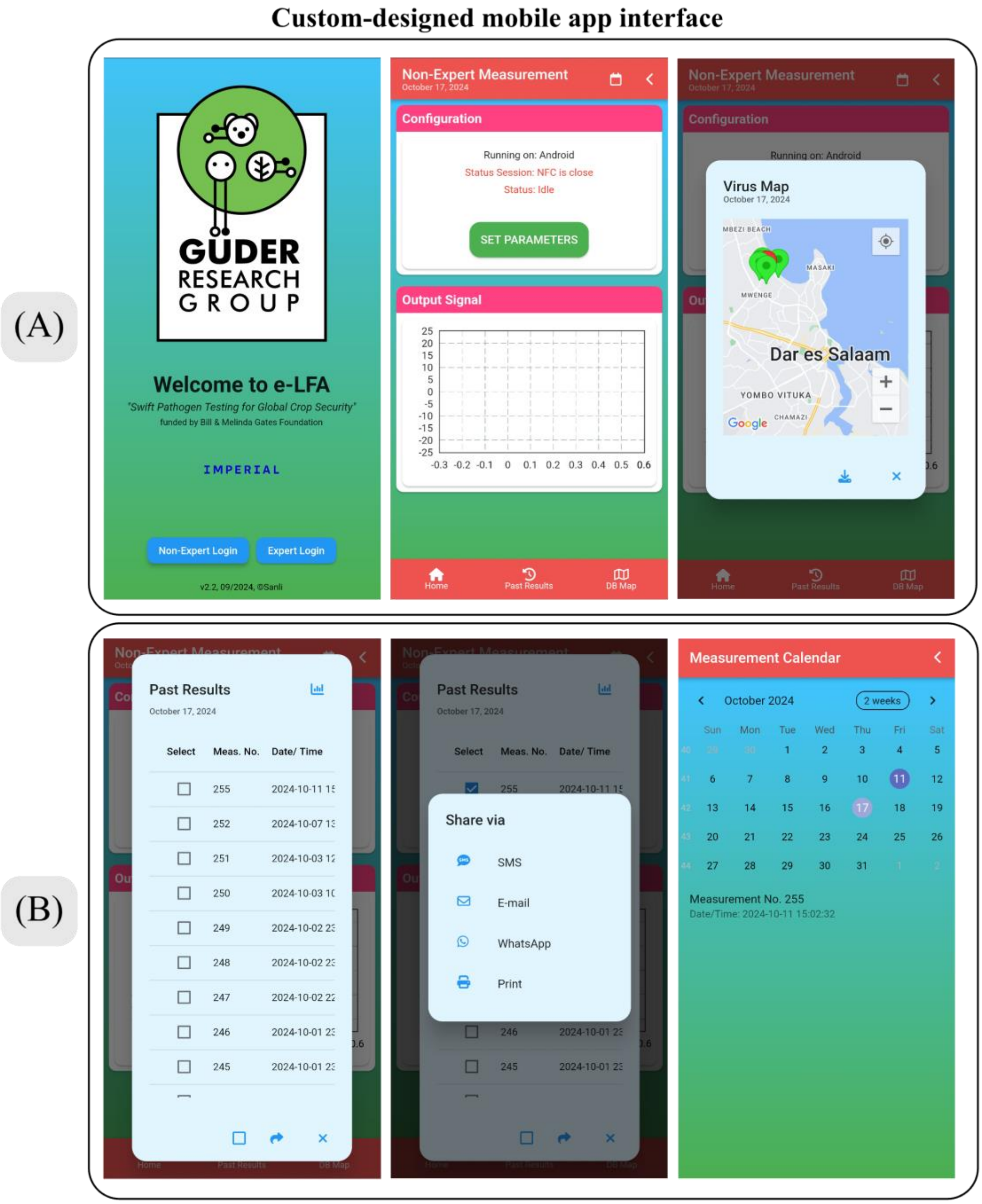


**Figure S4**: Screenshots of the custom-designed mobile app interface, developed with SIC4341 libraries in Flutter using the Dart programming language on Android Studio. (A) The configuration page allows users (expert and non-expert) to input settings and view a real-time virus map, which is automatically updated from a MongoDB database. This page provides essential controls for configuring device parameters, ensuring ease of use across different skill levels. (B) The measurement page presents a comprehensive list of all recorded measurements retrieved from both local and cloud-based databases, with options for data sharing and printing. A measurement calendar enables users to select specific dates to access and review individual measurements. This database system serves as a centralized repository for data sharing, encompassing experimental results, location-based information, and other relevant metrics, allowing seamless collaboration and broad accessibility across platforms.

**Electrodes characterization**. Scanning electron microscopy (SEM) of the gold-plated acupuncture needles revealed a smooth and uniform surface morphology along the shaft, with occasional microscopic imperfections and pits likely originating from the gold coating or underlying stainless-steel substrate. These minor surface defects are typical of commercially fabricated acupuncture needles and did not appear to disrupt the overall coating integrity. The absence of significant roughness or porosity suggests that the electrode surface is well-suited for reproducible electron transfer. No delamination or major inconsistencies were observed across multiple needles, supporting the use of these low-cost materials as reliable and consistent electrochemical interfaces.

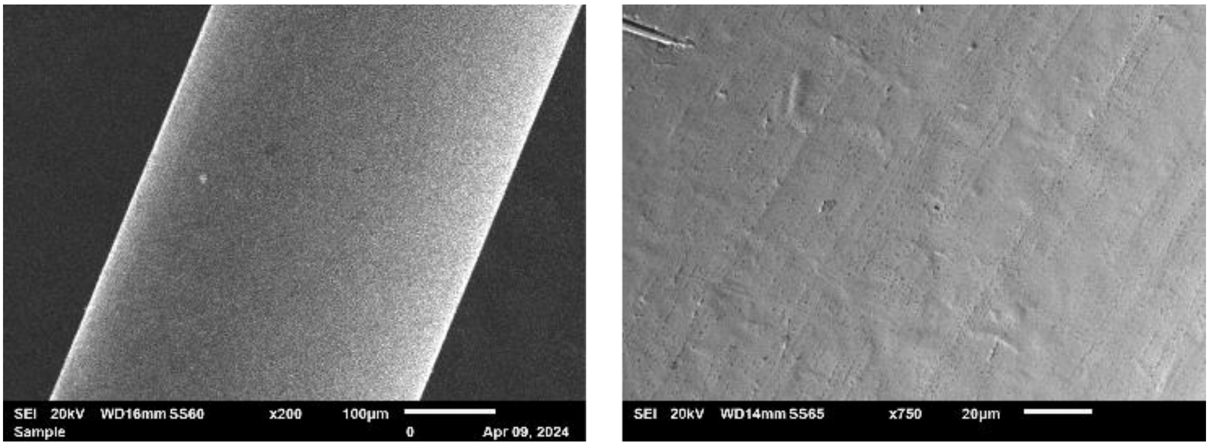


**Figure S5**: Scanning electron microscopies of the needles used as electrodes.

Cyclic voltammetry (CV) was carried out to characterize the electrochemical behavior of screen-printed gold electrodes (SPGEs) and the acupuncture-needle electrodes. A 2 mM solution of hexaammineruthenium(II/III) chloride (RuHex) in phosphate-buffered saline (PBS) was used as the redox probe. For the needle configuration, a silver-plated acupuncture needle served as the reference electrode, while gold-plated acupuncture needles were used as the working and counter electrodes; the needles were directly connected to a PalmSens 4 potentiostat. All electrodes (working, reference, and counter) were immersed 1 cm into the electrolyte solution. CVs were recorded at scan rates of 1000, 500, 250, 100, 50, 25, 10, and 5 mV $s^{-1}$ between -0.5 V and 0.1 V. We selected RuHex because, when using ferri/ferrocyanide under the same conditions, we consistently observed a triangular (clipped) oxidation peak and, especially at low scan rates, and a pronounced potential drift; RuHex provided stable, well-defined responses across the entire scan-rate range.

The needle-based electrodes exhibited a strong linear relationship between peak current and the square root of the scan rate, with $R^2 = 0.998$ for oxidation and $R^2 = 0.995$ for reduction, indicating highly reversible and diffusion-limited electron transfer. For comparison, commercial screen-printed gold electrodes (SPGEs 220AT, Metrohm-Drop Sens) tested under the same conditions showed similarly high correlation coefficients ($R^2 = 0.999$ for oxidation and $R^2 = 0.997$ for reduction) and nearly identical redox peak positions. These observations confirmed that both systems display near ideal electrochemical behavior with the $[Ru(NH_3)_6]^{3+/2+}$ probe. Using a diffusion coefficient of $7.1 \times 10^{-6}$ $cm^2 \cdot s^{-1}$ for $[Ru(NH_3)_6]^{3+}$, [2]we estimated the electroactive surface area for the Au needle electrodes from the slope of the Randles–Ševčík plots. The needle electrodes yielded a calculated electroactive area of 7.1 $mm^2$, while the SPGE electrodes had a lower electroactive area of 4.5 $mm^2$ although both types of electrodes had the same geometric area (12.5$mm^2$) in contact with the electrolyte solution. This discrepancy suggests that a smaller portion of the SPGE surface is electrochemically active, potentially due to the presence of non-conductive components in the ink, whereas metallic continuity of the needles likely provided a more accessible, electroactive surface for the redox reactions. Overall, the needle electrodes demonstrated superior electrochemical performance, making them the preferred choice for integration into the ELLA cassette.

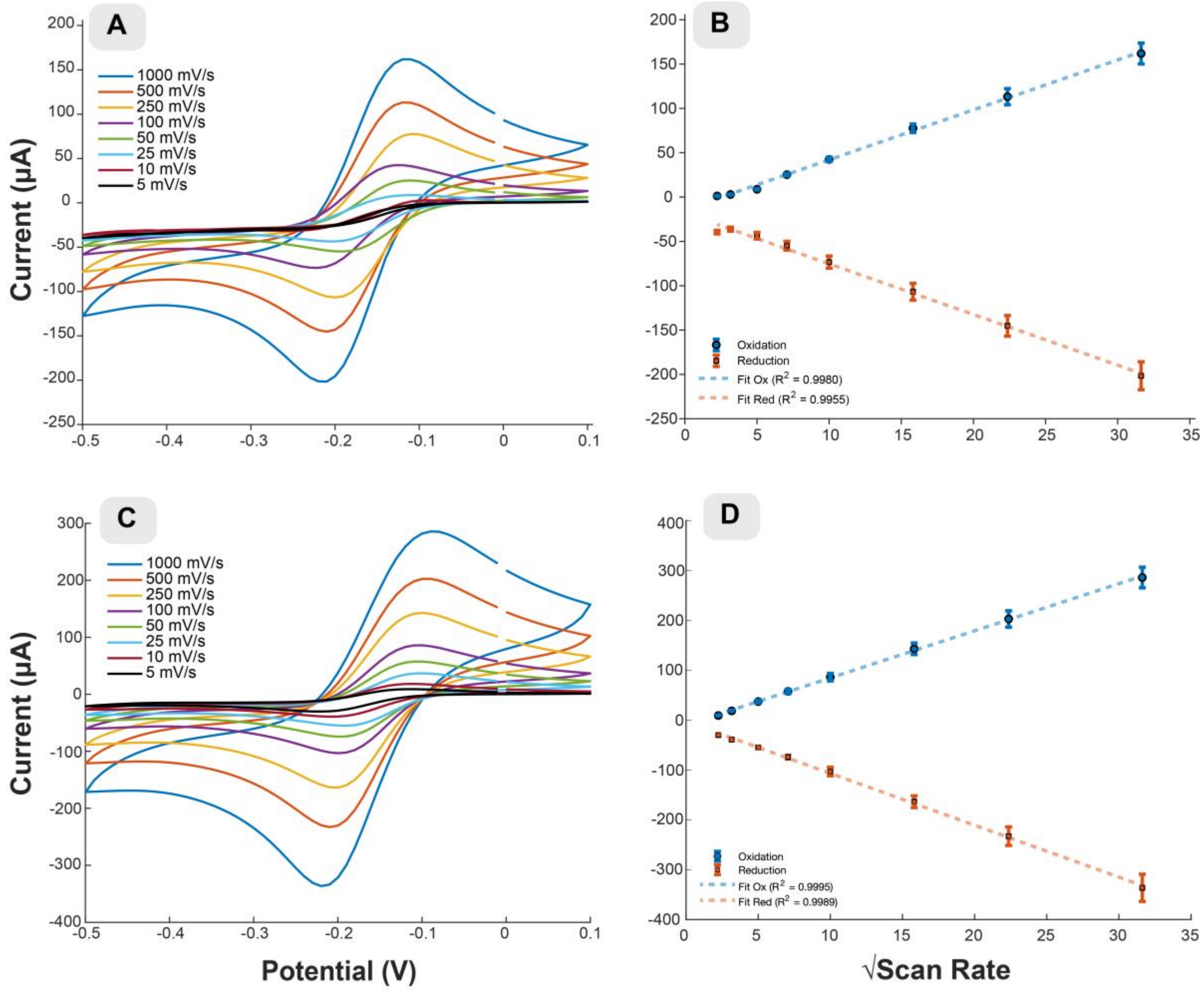


**Figure S6: (A)** Cyclic voltammograms of gold acupuncture needle electrodes recorded at scan rates of 5, 10, 25, 50, 100, 250, 500, and 1000 mV·s$^{-1}$ in 2 mM hexaammineruthenium(III)/(II) chloride dissolved in PBS (pH 7.4). The electrochemical cell consisted of gold-plated acupuncture needles as the working and counter electrodes, and a silver-plated acupuncture needle as the pseudo-reference electrode. All electrodes were immersed 1 cm into the electrolyte and connected to a benchtop potentiostat. **(B)** Corresponding peak anodic and cathodic currents plotted as a function of the square root of the scan rate, demonstrating linearity consistent with diffusion-controlled electrochemical behavior. **(C–D)** Identical measurements and analysis performed using commercial screen-printed gold electrodes (SPGEs), showing comparable voltammetric profiles and peak current dependence on scan rate.

**Platform comparison.** To evaluate the analytical performance of the custom-built PCB potentiostat relative to the commercial benchtop system (Autolab PGSTAT204, Metrohm), square-wave voltammetry (SWV) was performed in different concentrations of potassium ferri/ferrocyanide ($Fe(CN)_6^{3-/4-}$), dissolved in phosphate-buffered saline. For both instruments, identical three-electrode configurations were employed, ensuring the same geometric electrode area and electrolyte volume. The SWV parameters were: range from -0.8 to +0.8, amplitude 50mV, frequency 2.5 Hz. The peak current was extracted and averaged for each concentration. Calibration curves were generated (**Fig. S7**) by plotting the mean peak current against analyte concentration, with error bars representing the standard deviation (n = 3). The resulting datasets were fitted with a logarithmic regression models, yielding coefficients of determination ($R^2$) of 0.975 and 0.976 for the PCB and Autolab systems, respectively. The fitted equations were ip = 5.1403ln(x) + 10.1730 for the PCB and ip =5.1605ln(x) +1 1.8925, where ip is the SWV peak current (μA) and x the $Fe(CN)_6^{3-}/^{4-}$ concentration (mM). Both instruments exhibited a similar concentration-dependent response, characterized by a steep current increase at low concentrations followed by mild saturation above 2.5 mM, consistent with diffusion-limited behavior of the redox couple. The PCB potentiostat consistently produced currents approximately 17% lower in magnitude than the Autolab, attributable to differences in front-end amplification and current scaling; however, the sensitivity trend and non-linear shape were effectively identical. These results confirm that the PCB potentiostat reproduces the electrochemical response of the commercial instrument with fidelity, validating its use as an embedded readout unit for the ELLA platform.

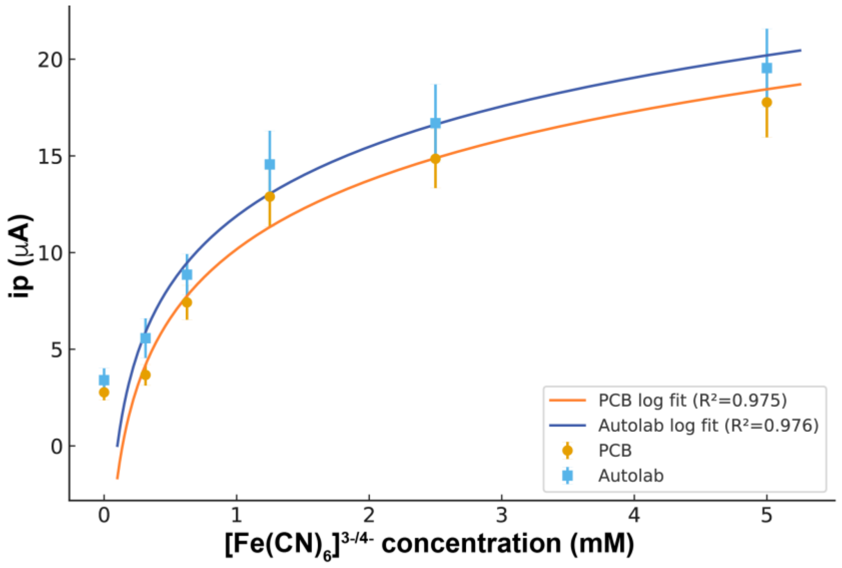


**Figure S7:** Comparison of calibration curves obtained using the custom PCB potentiostat and a commercial Autolab system. Square-wave voltammetry (SWV) peak currents recorded for increasing concentrations of ferri/ferrocyanide in PBS (0–5 mM) were fitted with a logarithmic function (potential range from -0.8 to +0.8, amplitude 50mV, frequency 2.5 Hz, n=3)

**Assembly process of test strips for ELLA.** The fabrication of the test strips involves preparing the nitrocellulose membrane and assembling key components, including the absorbent pad, nitrocellulose membrane, and the sample + AuNPs conjugate pad. Fig. S8 showcases both pre-test and post-test views, highlighting the appearance of a positive result.

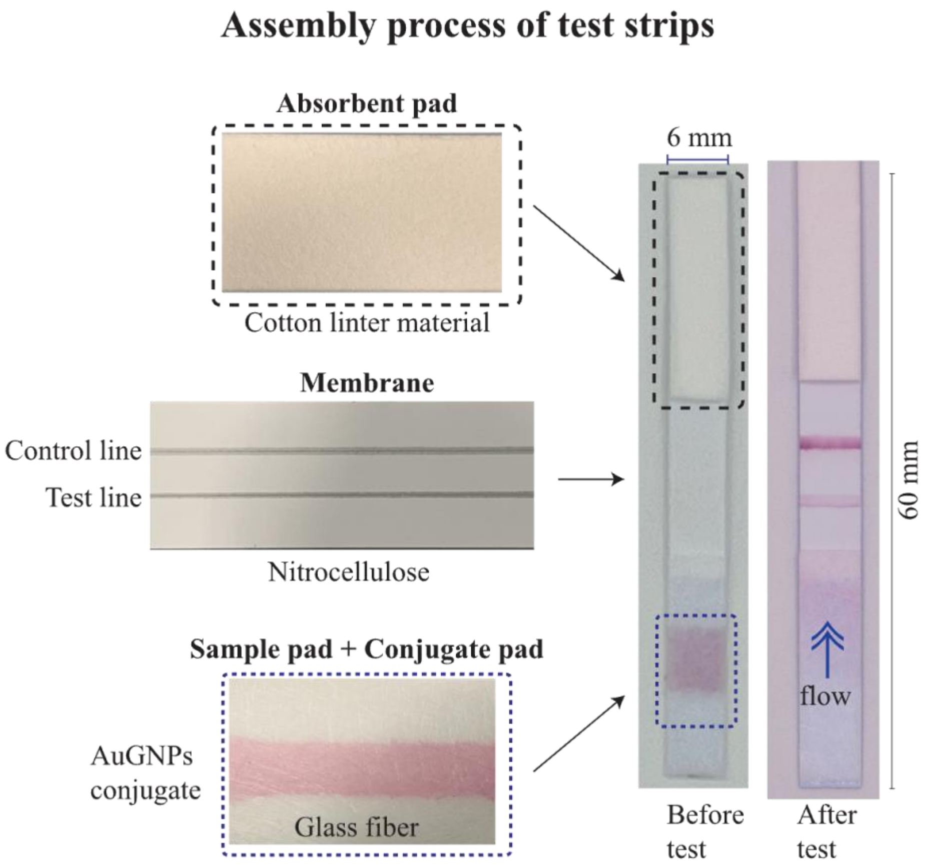


**Figure S8**: Assembly process of the test strips, including the preparation of the nitrocellulose membrane and the assembly of the test strip components: the absorbent pad, nitrocellulose membrane, and sample + AuNPs conjugate pad. Both pre-test and post-test views are shown, demonstrating the appearance of a positive result.

**Design of a 3D printed custom-made hole puncher device**. We designed a custom 3D-printed puncher that simultaneously aligns the nitrocellulose test strip with the PCB, which already carries the soldered needle electrodes (**Fig. S9**). In a single operation, the puncher perforates the strip and positions the electrodes through the membrane just after the test line, maintaining the structural integrity of the paper and ensuring precise, reproducible placement for subsequent measurements. The tip of the needle can be modified to ensure residue-free holes.

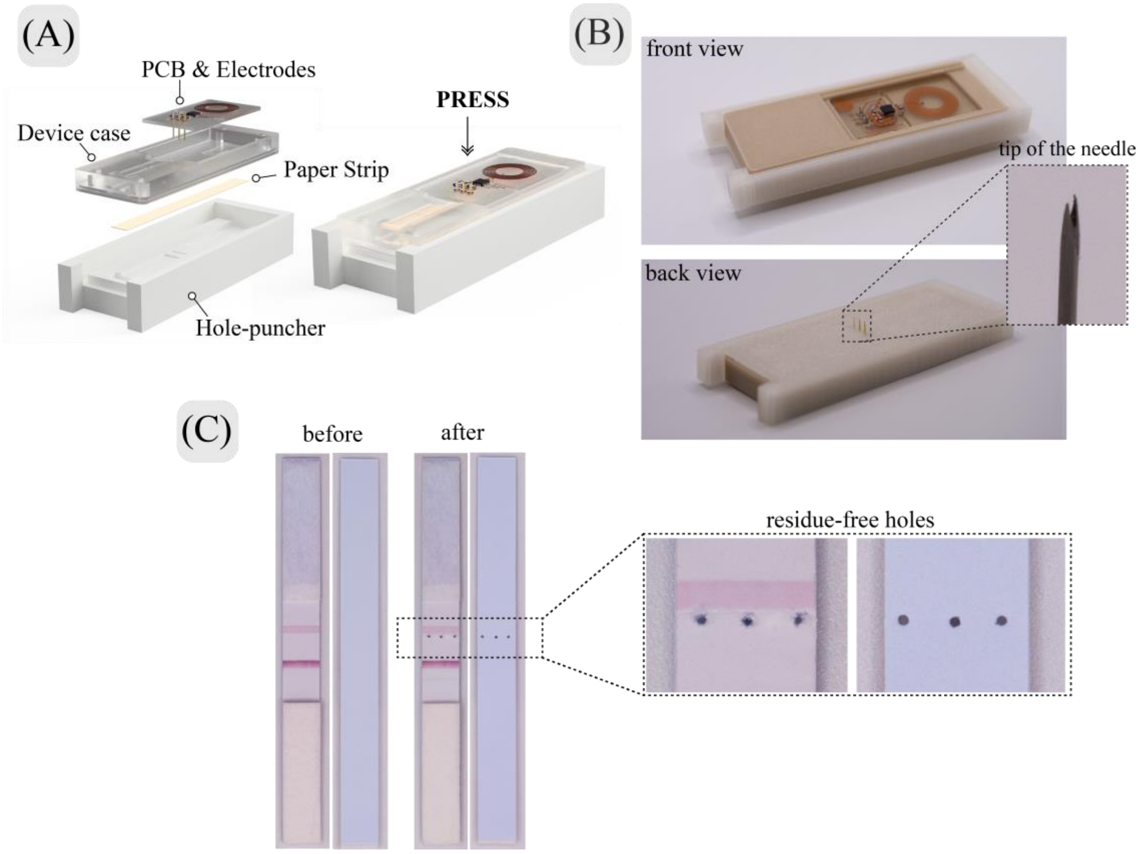


**Figure S9**: (A) 3D sketch and (B) photograph of the custom-designed hole puncher device, illustrating the integration of three curved-tip needles for precise perforation of nitrocellulose paper. (C) Front and back views of the punched sample demonstrate the device's capability to create consistent, residue-free holes, enhancing measurement reliability.

**Stability test.** To assess the shelf-life of the ELLA cassettes, the devices were stored at room temperature (25 °C ± 1), protected from light and accompanied by silica gel pouches to maintain low humidity levels. The stability study was conducted over a 5-month period, during which the ELLA cassettes were tested monthly using a positive control sample to monitor performance consistency. The results revealed a gradual decline in current, with current readings exhibiting a linear decrease, particularly noticeable from the second month of storage onwards. This decline suggests potential degradation of the active components or changes in the electrochemical properties over time. These findings highlight the importance of further optimization in the storage protocol or component stability to extend the effective shelf-life of the devices.

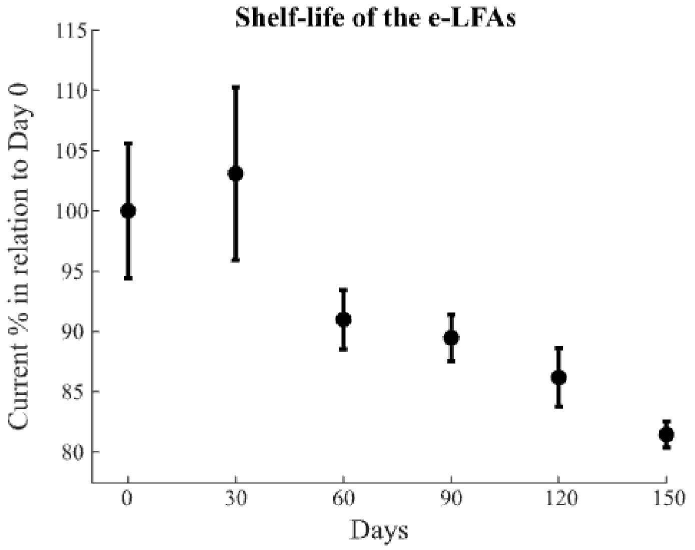


**Figure S10**: Shelf-life evaluation of ELLAs over a 5-month storage period. Devices were stored at 20°C, protected from light, and paired with silica pouches to maintain a dry environment. Performance was assessed monthly using a positive control sample to monitor functionality and stability over time.

**AI Classification of SWV.** SWV data were collected from 125 independent measurements. Each voltammogram was processed to extract both statistical and signal-shape features for machine learning classification. The following features were computed: maximum and minimum current, mean and standard deviation of current, area under the current–potential curve, maximum and minimum of first derivative (slope), skewness of the current distribution, peak potential, peak prominence, fast Fourier transform (FFT) descriptors, and polynomial fitting coefficients. Feature correlation analysis was performed to identify independent contributors. From the initial set, 13 features were found to provide non-redundant information. The most relevant among these included maximum/minimum current, skewness of the current shape, peak potential, peak width, FFT-derived descriptors, and polynomial coefficients. Feature importance was estimated by training ensemble models and ranking their contribution to classification accuracy (Fig. S10). Based on this analysis, the eight most informative features were selected for model development. Five supervised learning algorithms were evaluated: logistic regression, random forest, extreme gradient boosting (XGBoost), support vector machine (SVM with RBF kernel), and a multilayer perceptron (MLP, feed-forward neural network). For each model, hyperparameters were optimized through grid/random search where applicable. Model performance was assessed using repeated stratified k-fold cross-validation (k=5, 10 repeats), with 80% of data used for training and 20% held out for testing. All models achieved broadly comparable performance, with test set accuracies around 83% and ROC AUC values between 0.83–0.86. This performance was lower than that obtained using a simple thresholding approach on peak current alone, as reported elsewhere in this study. Nonetheless, the machine learning analysis demonstrates that SWV data contain rich, multidimensional information that can be leveraged for classification, and may yield further improvements with larger datasets or advanced feature engineering.

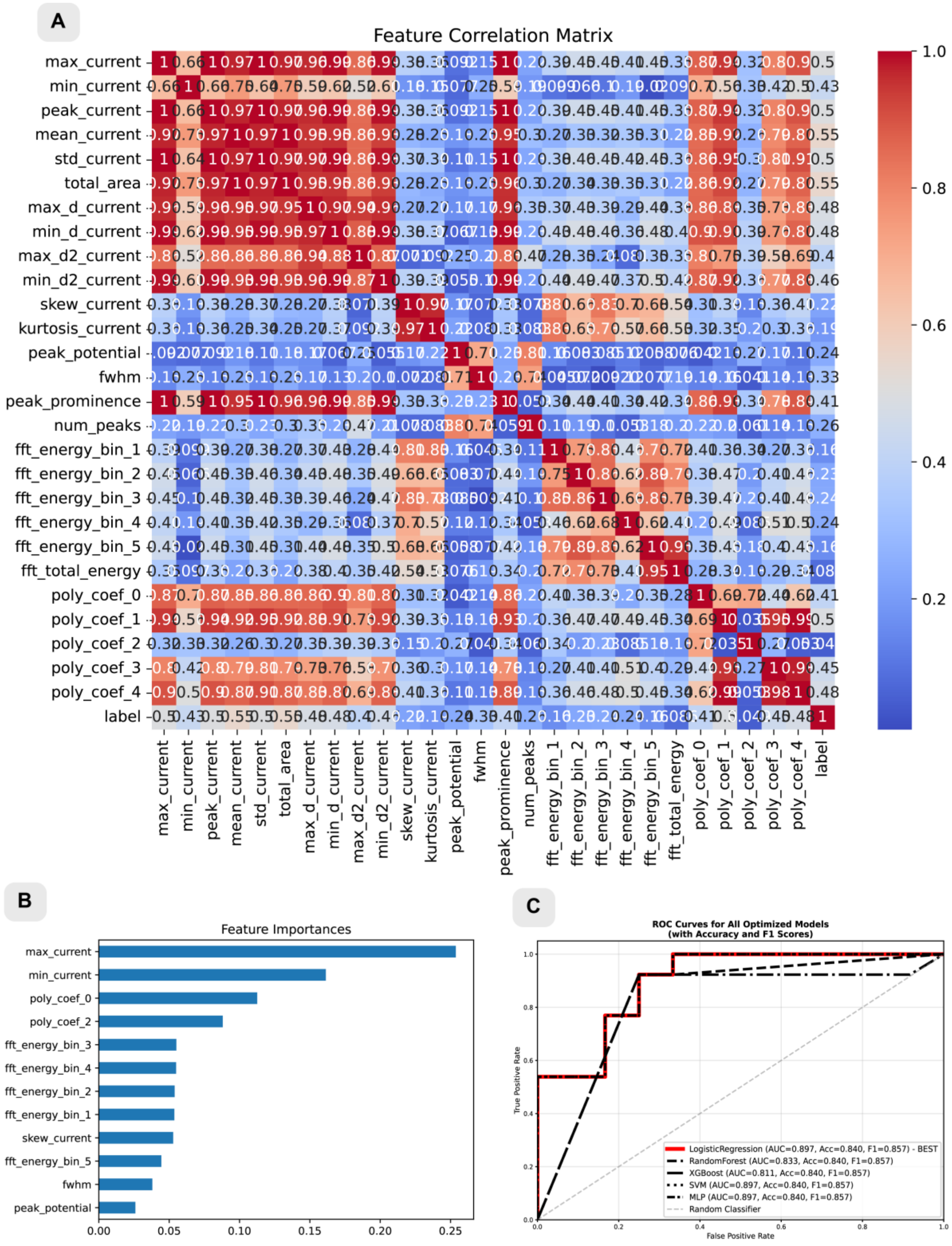


**Figure S11:** Feature evaluation and performance of machine learning models for CBSV detection. (**A**) Feature correlation matrix of the processed dataset (27 manually-derived features from SWV data), constructed from the absolute Pearson correlation coefficients among all extracted features. The heatmap highlights redundancy and potential multicollinearity, which were used to guide feature reduction by removing highly correlated variables

(> 0.9). (**B**) Feature importance analysis using a Random Forest classifier trained on the reduced dataset. Feature importance values were calculated from the mean decrease in Gini impurity, providing a relative ranking of each feature's contribution to model decision-making. The two most-important features were selected for model training as increasing the number had no effect or deteriorated model performance. (**C**) Receiver operating characteristic (ROC) curves generated for all optimized models following stratified train–test splitting (80/20). Hyperparameter optimization was performed on five models (logistic regression, random forest, XGBoost, support vector machine, and multilayer perceptron) using grid search with stratified 5-fold cross-validation on the training set to identify the parameter set yielding the best mean performance. The final models were then retrained with the selected hyperparameters and evaluated on the test set. ROC curves are shown alongside accuracy and F1 scores for each model.

**ELISA and RT-qPCR tests**. The ELISA kit for detecting CBSV was obtained from the Leibniz Institute DSMZ (Germany) and was conducted according to the recommended guidelines. Briefly, it involves a triple antibody sandwich format. The process begins with coating 96-well plates with purified IgG against CBSV (AS-1153) diluted in coating buffer, followed by incubation and washing. A blocking step with 2% skim milk in PBS-Tween is performed to prevent nonspecific reactions. Leaf samples are ground in sample extraction buffer (1:10 w/w ratio) and incubated overnight at 4°C. After washing, monoclonal antibodies (MAb AS-1153/1) are added, followed by incubation and another wash. A secondary anti-mouse antibody conjugated with alkaline phosphatase (RAM-AP) is then added, followed by incubation and washing. Finally, a substrate solution of p-nitrophenyl phosphate is added, incubated and the results are measured by optical density at 405 nm. Positive results are indicated by absorbance values three times higher than the means of negative controls.

Reverse transcription-quantitative polymerase chain reaction (RT-qPCR) was performed using a duplex TaqMan assay employing primers and probes designed to simultaneously detect CBSV and UCBSV (Shirima et al.; unpublished data). Portions (lobes) from the cassava leaf samples that were selected for the ELLA and ELISA tests were attached to blank newsprint papers arranged in layers of two to three and left to dry at ambient temperature. Approximately 30 mg of the dried leaf was subjected to RNA extraction using a modified cetyltrimethylammonium bromide method (Maruthi et al. 2002). The recovered RNA pellet was resuspended in nuclease-free PCR-grade water immediately or stored at –20 °C for later use. In setting up the RT-qPCR reaction, 1 µL of RNA extract was used in a 25 µL reaction volume. The duplex TaqMan assay mix contained: 1× PCR buffer, 5.5 mM MgCl2, 0.5 mM dNTPs [New England Biolabs], 0.3 µM CBSV forward primer, 0.3 µM CBSV reverse primer, 0.1 µM CBSV probe, 0.4 µM UCBSV forward primer, 0.4 µM UCBSV reverse primer, 0.2 µM UCBSV probe (primers and probes were synthesized by IDT DNA Technologies, Inc.), 200 nM ROX, 1 unit of Taq DNA polymerase and 17 units of M-MLV reverse transcriptase (all PCR reagents except dNTPS were obtained from Thermo Fisher Scientific). The reaction was run on the AriaMx Q-PCR System, Agilent Technologies: Santa Clara, CA 95051 United States. Thermal cycling was set as 48 °C for 30 minutes, with initial denaturation at 95 °C for 10 minutes followed by 40 cycles of 95 °C for 15 seconds, 52 °C for 20 seconds, 55 °C 15 seconds and 60 °C (optical data collection) for 40 seconds. The quantification cycle values (Cq) were generated using the onboard AriaMx Software version 2.1.

**Table S1 –** Comparison of ELLA current with standard laboratory tests for CBSD detection.

| Sample No. | Variety | CBSD stage | ELLA peak current (µA)* | qPCR quantification cycle (Cq)** | ELISA absorbance*** |
|---|---|---|---|---|---|
| 1 | Albert | 1 | 2.65 | 0.0 | 0.451 |
| 2 | Albert | 3 | 1.52 | 0.0 | 1.378 |
| 3 | Albert | 3 | 0.99 | 0.0 | 0.667 |
| 4 | Albert | 1 | 1.18 | 0.0 | 0.441 |
| 5 | Albert | 3 | 1.19 | 0.0 | 0.384 |
| 6 | Albert | 3 | 4.52 | 0.0 | 0.435 |
| 7 | Albert | CMD | 0.76 | 0.0 | 0.867 |
| 8 | Albert | 1 | 0.86 | 0.0 | 0.637 |
| 9 | Albert | 3 | 4.32 | 32.3 | 1.144 |
| 10 | Albert | 3 | 5.15 | 13.1 | 1.162 |
| 11 | Albert | 1 | 2.77 | 0.0 | 0.395 |
| 12 | Albert | 3 | 0.93 | 0.0 | 0.436 |

| | | | | | |
|---|---|---|---|---|---|
| 13 | Albert | 3 | 4.2 | 14.2 | 1.608 |
| 14 | Albert | CMD | 1.73 | 0.0 | 0.687 |
| 15 | Albert | 1 | 2.48 | 0.0 | 0.728 |
| 16 | Albert | 3 | 5.23 | 12.7 | 1.214 |
| 17 | Albert | 3 | 3.41 | 13.5 | 0.563 |
| 18 | Kiroba | 1 | 2.52 | 0.0 | 0.535 |
| 19 | Kiroba | 3 | 12.58 | 31.8 | 1.798 |
| 20 | Kiroba | 3 | 5.28 | 15.3 | 0.857 |
| 21 | Kiroba | 1 | 1.1 | 0.0 | 0.714 |
| 22 | Kiroba | 3 | 5.24 | 0.0 | 1.256 |
| 23 | Kiroba | 3 | 1.81 | 0.0 | 0.57 |
| 24 | Kiroba | 1 | 0.54 | 0.0 | 0.381 |
| 25 | Kiroba | 2 | 2.73 | 0.0 | 0.375 |
| 26 | Kiroba | 1 | 0.74 | 0.0 | 0.412 |
| 27 | Kiroba | 2 | 3.25 | 0.0 | 0.642 |
| 28 | Kiroba | 1 | 1.9 | 0.0 | 0.45 |
| 29 | Kiroba | 3 | 8.44 | 0.0 | 1.066 |
| 30 | Kiroba | 3 | 4.69 | 0.0 | 0.728 |
| 31 | Nyoku-Agbeli | 1 | 3.25 | 0.0 | 0.686 |
| 32 | Nyoku-Agbeli | 3 | 11.85 | 13.3 | 0.943 |
| 33 | Nyoku-Agbeli | 3 | 6.84 | 0.0 | 0.938 |
| 34 | Nyoku-Agbeli | 1 | 1.04 | 0.0 | 0.619 |
| 35 | Nyoku-Agbeli | 3 | 10.68 | 15.1 | 0.845 |
| 36 | Nyoku-Agbeli | 3 | 4.52 | 0.0 | 0.937 |
| 37 | Nyoku-Agbeli | 1 | 0.57 | 0.0 | 0.539 |
| 38 | Nyoku-Agbeli | CMD | 1.02 | 0.0 | 0.698 |
| 39 | Nyoku-Agbeli | 2 | 12.23 | 11.9 | 1.575 |
| 40 | Nyoku-Agbeli | 2 | 2.42 | 0.0 | 0.353 |
| 41 | Fufu-Bankye | 1 | 4.28 | 13.1 | 1.015 |
| 42 | Fufu-Bankye | 2 | 5.65 | 20.1 | 1.543 |
| 43 | Fufu-Bankye | 1 | 1.14 | 0.0 | 0.472 |
| 44 | Fufu-Bankye | 3 | 14.22 | 4.3 | 0.936 |
| 45 | Fufu-Bankye | 3 | 13.16 | 10.8 | 1.081 |
| 46 | UCC2001-053 | 1 | 3.5 | 0.0 | 0.461 |
| 47 | UCC2001-053 | 3 | 4.76 | 12.7 | 1.638 |
| 48 | UCC2001-053 | 3 | 10.94 | 0.0 | 0.526 |
| 49 | UCC2001-053 | 1 | 3.16 | 0.0 | 0.464 |
| 50 | UCC2001-053 | 3 | 7.24 | 18.2 | 1.506 |
| 51 | UCC2001-053 | 3 | 8.55 | 11.4 | 0.843 |
| 52 | Albert (Old) | 1 | 1.54 | 22.6 | 0.454 |

| | | | | | |
|---|---|---|---|---|---|
| 53 | Albert (Old) | 4 | 7.63 | 15.9 | 1.088 |
| 54 | Albert (Old) | 4 | 5.18 | 13.0 | 1.815 |
| 55 | Albert (Old) | 4 | 29.41 | 15.6 | 1.864 |
| 56 | Albert (Old) | 1 | 8.03 | 21.7 | 1.653 |
| 57 | Albert (Old) | 4 | 1.74 | 16.7 | 0.711 |
| 58 | Albert (Old) | 4 | 2.21 | 13.2 | 0.474 |
| 59 | Albert (Old) | 4 | 4.22 | 13.3 | 1.777 |
| 60 | Kiroba (Old) | 1 | 3.62 | 18.6 | 0.911 |
| 61 | Kiroba (Old) | 3 | 9.29 | 14.7 | 1.039 |
| 62 | Kiroba (Old) | 3 | 13.46 | 14.5 | 1.435 |
| 63 | Kiroba (Old) | 1 | 6.49 | 23.1 | 1.322 |
| 64 | Kiroba (Old) | 3 | 8.79 | 17.8 | 1.761 |
| 65 | Kiroba (Old) | 3 | 11.83 | 19.3 | 0.962 |
| 66 | Kiroba (Old) | 1 | 7.01 | 20.2 | 1.564 |
| 67 | Kiroba (Old) | 3 | 8.68 | 19.0 | 1.04 |
| 68 | Kiroba (Old) | 3 | 15.11 | 19.0 | 0.9 |
| 69 | Kiroba (Old) | 1 | 8.64 | 21.5 | 0.92 |
| 70 | Kiroba (Old) | 3 | 13.65 | 19.8 | 1.737 |
| 71 | Kiroba (Old) | 3 | 13.47 | 19.7 | 1.504 |
| 72 | Albert (Old) | 1 | 1.39 | 0.0 | 0.433 |
| 73 | Albert (Old) | 3 | 2.4 | 0.0 | 0.491 |
| 74 | Albert (Old) | 3 | 11.25 | 0.0 | 0.946 |
| 75 | Albert (Old) | 1 | 3.35 | 0.0 | 0.6 |
| 76 | Albert (Old) | 3 | 0.73 | 0.0 | 0.743 |
| 77 | Albert (Old) | 3 | 1.73 | 0.0 | 0.534 |
| 78 | Albert (Old) | 1 | 0.83 | 0.0 | 0.413 |
| 79 | Albert (Old) | 4 | 7.87 | 0.0 | 0.955 |
| 80 | Albert (Old) | 4 | 3.33 | 0.0 | 0.698 |
| 81 | Albert (Old) | 4 | 0.78 | 0.0 | 0.633 |
| 82 | Albert | 1 | 2.76 | 0.0 | 0.462 |
| 83 | Albert | 2 | 2.34 | 0.0 | 0.417 |
| 84 | Albert | 2 | 2.12 | 0.0 | 0.458 |
| 85 | Albert | 2 | 8.07 | 17.9 | 1.18 |
| 86 | 031 | 1 | 2.11 | 0.0 | 0.657 |
| 87 | 031 | 5 | 1.98 | 0.0 | 0.401 |
| 88 | 031 | 5 | 10.67 | 0.0 | 1.472 |
| 89 | 031 | 5 | 14.15 | 17.0 | 1.591 |
| 90 | 031 | 5 | 22.7 | 15.6 | 0.979 |
| 91 | 031 | 1 | 3.84 | 25.1 | 0.568 |
| 92 | 031 | 5 | 9.35 | 22.3 | 0.881 |

| | | | | | |
|---|---|---|---|---|---|
| 93 | 031 | 5 | 17.01 | 17.2 | 0.91 |
| 94 | 031 | 5 | 31.79 | 16.5 | 1.666 |
| 95 | 031 | 5 | 30.46 | 17.2 | 1.187 |
| 96 | 031 | 1 | 1.42 | 0.0 | 0.503 |
| 97 | 031 | 3 | 2.27 | 0.0 | 0.674 |
| 98 | 031 | 3 | 21.53 | 15.6 | 1.027 |
| 99 | UKG 09-152 | 1 | 3.2 | 0.0 | 0.338 |
| 100 | UKG 09-152 | 4 | 15.5 | 23.3 | 1.166 |
| 101 | UKG 09-152 | 4 | 24.44 | 19.9 | 1.617 |
| 102 | UKG 09-152 | 4 | 17.01 | 15.8 | 1.98 |
| 103 | UKG 09-152 | 1 | 13.8 | 24.8 | 1.439 |
| 104 | UKG 09-152 | 4 | 16.7 | 18.0 | 1.622 |
| 105 | UKG 09-152 | 4 | 12.37 | 15.9 | 1.234 |
| 106 | UKG 09-152 | 4 | 15.04 | 17.1 | 1.475 |
| 107 | UKG 09-152 | 1 | 1.86 | 0.0 | 0.783 |
| 108 | UKG 09-152 | 3 | 10.05 | 17.5 | 0.911 |
| 109 | UKG 09-152 | 3 | 11.91 | 17.1 | 1.122 |
| 110 | Albert | 1 | 2.61 | 0.0 | 0.567 |
| 111 | Albert | 2 | 2.91 | 22.8 | 0.525 |
| 112 | Albert | 1 | 1.09 | 0.0 | 0.694 |
| 113 | Albert | 2 | 5.53 | 21.0 | 0.602 |
| 114 | Albert | 1 | 9.48 | 0.0 | 0.752 |
| 115 | Albert | 2 | 3.86 | 22.2 | 0.644 |
| 116 | Albert | 2 | 4.65 | 17.9 | 1.122 |
| 117 | Kiroba | 3 | 9.76 | 17.0 | 1.823 |
| 118 | Albert | 3 | 17.81 | 16.8 | 1.672 |
| 119 | TME 14 | 3 | 10.66 | 16.8 | 1.342 |
| 120 | TME 204 | 3 | 7.54 | 16.2 | 1.156 |
| 121 | Albert | 1 | 1.19 | 0.0 | 0.547 |
| 122 | Kiroba | 1 | 4.03 | 0.0 | 0.634 |
| 123 | TME 204 | 3 | 5.14 | 17.1 | 0.823 |
| 124 | Albert | 3 | 6.29 | 17.3 | 1.042 |
| 125 | Albert | 1 | 1.05 | 0.0 | 0.531 |
| PC100% | - | - | 11.04 | 10.1 | 2.341 |
| PC50% | - | - | 10.27 | - | 1.230 |
| PC25% | - | - | 9.01 | - | 0.764 |
| PC10% | - | - | 6.55 | - | 0.341 |
| PC5% | - | - | 4.58 | - | 0.324 |
| NC | - | - | 2.51 | 0.0 | 0.250 |

*Current values > 4μA are considered positive for CBSD; **Cq values > 0 are considered positive for CBSD;

***Absorbance values > 0.75 are considered positive for CBSD (3x NC signal); PC = positive control; NC = Negative control; CMD = Plant infected with Cassava Mosaic Disease

**Table S2** – Cost of materials for one fully assembled ELLA platform.

| **Part** | **Cost* (USD)** |
|---|---|
| Antibodies | 0.40 |
| Nanoparticles | 0.10 |
| Nitrocellulose | 0.08 |
| Pads and backing card | 0.04 |
| 3D printed cassette | 0.03 |
| Electrodes | 0.03 |
| SIC4341 Silicon Craft Chip | 0.20 |
| Capacitors | 0.01 |
| PCBs | 0.01 |
| **Total** | **US$ 0.90** |

*Prices are converted from GBP using the prevailing exchange rate, subject to change. SIC4341 chips were obtained from Silicon Craft PLC and the rest of the electronic components were purchased from Mouser Electronics.

### References

1. Texas Instruments. Coil Designer. https://webench.ti.com/wb5/LDC/#/spirals (2024).
2. Wang, Y., Limon-Petersen, J. G. & Compton, R. G. Measurement of the diffusion coefficients of [Ru(NH3)6]3+ and [Ru(NH3)6]2+ in aqueous solution using microelectrode double potential step chronoamperometry. *J. Electroanal. Chem.* **652**, 13–17 (2011).
